# High functional diversity among *Nitrospira* populations that dominate rotating biological contactor microbial communities in a municipal wastewater treatment plant

**DOI:** 10.1101/529826

**Authors:** Emilie Spasov, Jackson M. Tsuji, Laura A. Hug, Andrew C. Doxey, Laura A. Sauder, Wayne J. Parker, Josh D. Neufeld

## Abstract

Nitrification, the oxidation of ammonia to nitrate via nitrite, is an important process in municipal wastewater treatment plants (WWTPs). Members of the *Nitrospira* genus that contribute to complete ammonia oxidation (comammox) have only recently been discovered and their relevance to engineered water treatment systems is poorly understood. This study investigated distributions of *Nitrospira*, ammonia-oxidizing archaea (AOA), and ammonia-oxidizing bacteria (AOB) in biofilm samples collected from tertiary rotating biological contactors (RBCs) of a municipal WWTP in Guelph, Ontario, Canada. Using quantitative PCR (qPCR), 16S rRNA gene sequencing, and metagenomics, our results demonstrate that *Nitrospira* species strongly dominate RBC biofilm samples and that comammox *Nitrospira* outnumber all other nitrifiers. Genome bins recovered from assembled metagenomes reveal multiple populations of comammox *Nitrospira* with distinct spatial and temporal distributions, including several taxa that are distinct from previously characterized *Nitrospira* members. Diverse functional profiles imply a high level of niche heterogeneity among comammox *Nitrospira*, in contrast to the sole detected AOA representative that was previously cultivated and characterized from the same RBC biofilm. Our metagenome bins also reveal two cyanase-encoding populations of comammox *Nitrospira*, suggesting an ability to degrade cyanate, which has not been shown previously for *Nitrospira* that are not strict nitrite oxidizers. This study demonstrates the importance of RBCs as model systems for continued investigation of environmental factors that control the distributions and activities of AOB, AOA, comammox *Nitrospira*, and other nitrite oxidizers.

## Introduction

Municipal wastewater contains ammonium that is removed by wastewater treatment plants (WWTPs) to prevent eutrophication, oxygen depletion, and toxicity to aquatic animals in receiving waters. Nitrification involves the sequential oxidation of ammonia to nitrate, via nitrite, and these two enzymatic steps were historically thought to be mediated by distinct microorganisms, with aerobic ammonia oxidation conducted by ammonia-oxidizing bacteria (AOB) or ammonia-oxidizing archaea (AOA), and nitrite oxidation catalyzed by nitrite-oxidizing bacteria (NOB) [1–3]. Microorganisms capable of oxidizing both ammonia and nitrite via complete ammonia oxidation (comammox) were predicted over a decade ago to be slow growing and to inhabit biofilms exposed to relatively low ammonium concentrations [4]. These predictions were confirmed by the discovery of *Nitrospira* members capable of catalyzing comammox [5, 6], and all known comammox *Nitrospira* belong to lineage II of the genus *Nitrospira*. Although two major clades of comammox *Nitrospira* have been described (i.e., clades A and B), based on ammonia monooxygenase (*amoA*) gene phylogeny, all enriched and cultivated species of comammox *Nitrospira* belong to clade A [5, 6].

Compared to AOA and AOB, very little is known about the abundance and diversity of comammox *Nitrospira* in engineered aquatic environments. Consistent with a low ammonium niche, comammox *Nitrospira* have been detected in drinking water systems [7–14]. First identified from water treatment system metagenome sequences [5, 6], most wastewater-associated comammox *Nitrospira* belong to clade A [5, 7, 12, 14–18], albeit with abundances generally lower than those reported for AOA and AOB [7, 18–21]. Nonetheless, *amoA* gene abundances of comammox *Nitrospira* may outnumber those of AOB in activated sludge samples in some WWTPs [12, 18, 22], and comammox *Nitrospira* have been enriched in wastewater treatment reactors with low dissolved oxygen conditions [15, 17]. Additionally, the high abundance of *amoA* transcripts from comammox *Nitrospira* detected in activated sludge suggests an active role in nitrification [23, 24]. Previous studies examining comammox *Nitrospira* in WWTPs focused primarily on activated sludge secondary treatment systems and sequencing batch reactors [12, 15–23, 25–28]. To our knowledge, no studies have investigated comammox *Nitrospira* in biofilm-based tertiary wastewater treatment systems.

The current study examined full scale tertiary rotating biological contactors (RBCs; Figure 1A) treating municipal waste from ∼132,000 residents of Guelph, Ontario, Canada. Comprising a total biofilm surface area of 440,000 m^2^ and processing ∼53,000 m^3^ of wastewater daily, the RBCs are organized into four “trains” that are each composed of eight individual RBC “stages” (Figure 1B). The RBC-associated AOA and AOB communities were characterized previously, demonstrating that the sole detectable AOA population increased in abundance along the RBC flowpath [29]. This AOA species was enriched, characterized, and named *Candidatus* Nitrosocosmicus exaquare [30]. However, inconsistent *in situ* activity data obtained using differential inhibitors for AOB, which showed differing inhibition patterns for AOB with two different AOB inhibitors, as well as an abundance of MAR-positive *Nitrospira* cells, as viewed by fluorescence *in situ* hybridization microautoradiography (FISH-MAR), raised the possibility that some of the detected *Nitrospira* may contribute to comammox [30]. Due to the predicted low ammonium niche and biofilm growth of comammox bacteria [4, 5], we hypothesized that comammox *Nitrospira* would dominate the RBC biofilm and that the relative abundance of comammox *Nitrospira* would increase as ammonium concentrations decrease along the RBC flowpath, as demonstrated previously for *Ca.* N. exaquare [29]. To test these hypotheses, we assessed the relative abundance, diversity, and temporal stability of comammox *Nitrospira*, in relation to AOA and other AOB, throughout the tertiary treatment system, using a combination of quantitative PCR (qPCR), 16S rRNA gene sequencing, and metagenomics.

**Figure 1.**
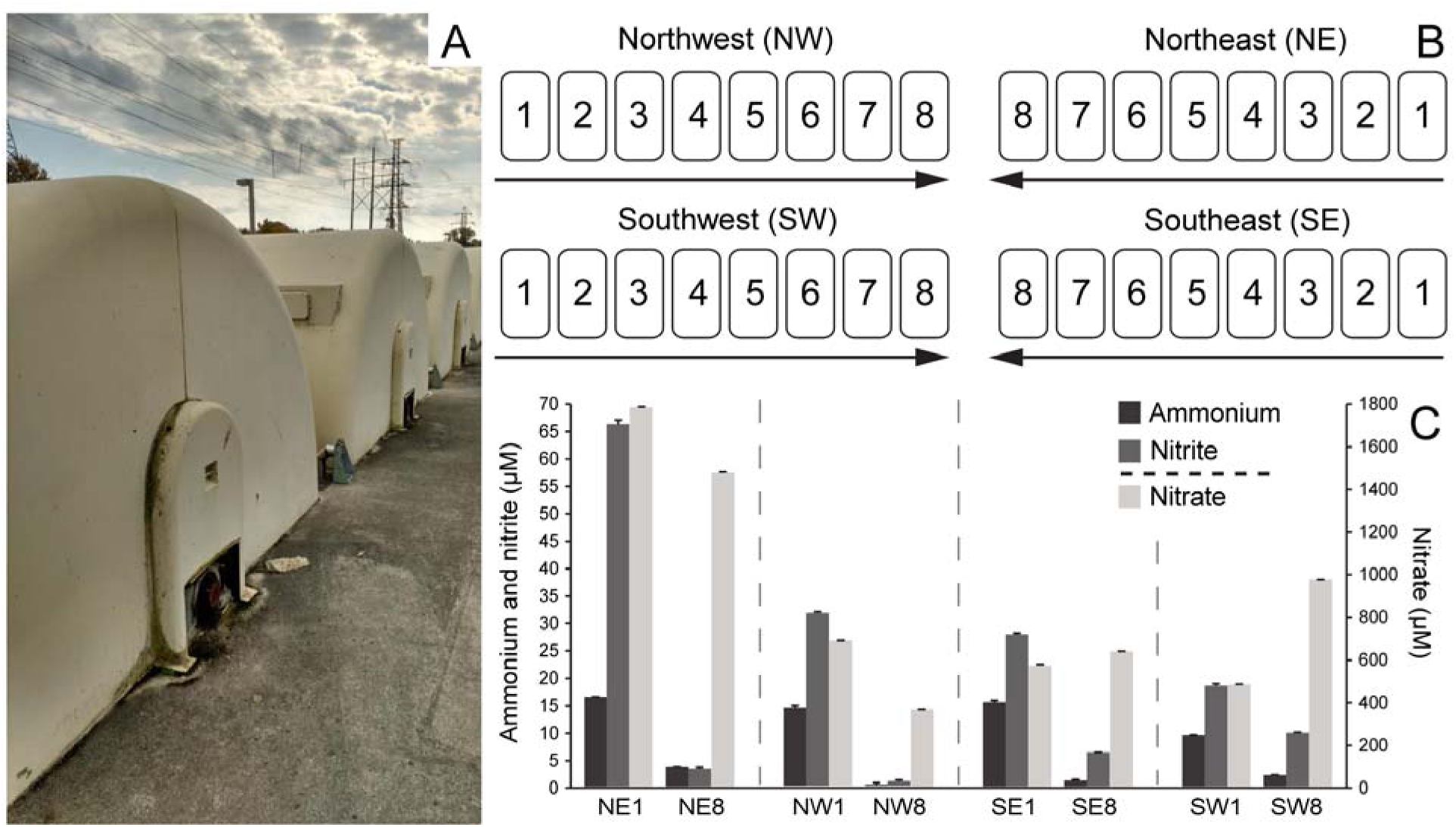
Rotating biological contactors (RBCs) of the Guelph wastewater treatment plant (WWTP). (A) External view of the RBCs and (B) schematic of the RBC trains. Arrows indicate the direction of water flow. (C) Ammonium (NH_4_^+^-N), nitrite (NO_2_-N), and nitrate (NO_3_-N) concentrations in RBC influent water sampled at the same time as biofilm collection in October 2016. Error bars indicate standard deviation of technical duplicates. NE, Northeast train; NW, Northwest train; SE, Southeast train; SW, Southwest train.

## Materials and Methods

### Sampling

Using ethanol-cleaned spatulas, biofilm was sampled from the RBC trains in Guelph, Ontario, Canada (Figure 1A, Figure 1B). Samples were stored on dry ice until delivered to the lab and were kept at −70°C until analyzed. All RBCs from all four trains were sampled in October 2016, except for RBC 1 of the southeast (SE) train and RBC 2 of the southwest (SW) train, which were not operational at the time of sampling (Table S1). We included biofilm samples that were collected previously for temporal comparison [29]. These reference samples were collected in February, June, and September 2010 from RBCs 1 and 8 of the northeast (NE) train (Table S1). Water samples were collected in 2016 from RBC 1 and RBC 8, kept on ice, and then frozen at −20°C until chemical analysis.

### Water chemistry

Ammonium was measured fluorometrically using orthophthaldialdehyde (OPA) reagent [31] as described previously [32] and nitrite/nitrate was assessed colorimetrically with the Griess reagent [33]. Measurements represent total ammonium as nitrogen, and nitrite and nitrate as nitrogen; details are described in the supplemental methods. Based on data obtained from WWTP operators, monthly and yearly ammonium concentrations (measured as total ammonia as nitrogen) of secondary effluent (i.e., RBC influent) and water from the southeast RBC train were compiled for the years 2010 to 2017. Plant influent and effluent data were retrieved from Guelph WWTP annual reports for 2010 to 2017 (https://guelph.ca/living/environment/water/wastewater/).

### DNA extractions

Extractions were performed from 0.25 g (wet weight) of biofilm with the PowerSoil DNA Isolation Kit (Mo Bio, Carlsbad, CA, USA) as described previously [30]. Total isolated DNA was visualized on a 1% agarose gel and quantified using the NanoDrop 2000 (Thermo Scientific, Waltham, MA, USA) and Qubit dsDNA HS Assay kit (Thermo Scientific).

Extracted DNA from RBC 1 and 8 biofilm samples of the NE train from 2010 was used for quantitative PCR (qPCR) and metagenome sequencing (Table S1). The 2010 biofilm samples were those collected and analyzed by qPCR previously [29], but DNA extractions were repeated while processing the 2016 samples. The DNA extracts from all RBCs within the four trains sampled in 2016 were analyzed with qPCR, whereas DNA from RBCs 1 and 8 alone were used for 16S rRNA gene and metagenome sequencing (Table S1).

### Quantitative PCR

Thaumarchaeotal and bacterial 16S rRNA genes were quantified using the primers 771F/957R [34] and 341F/518R [35], respectively (Table S2). Quantification of AOB *amoA* genes was carried out using the primers amoA1F/amoA2R [36] (Table S2). Comammox *Nitrospira* clade A and clade B *amoA* genes were amplified using equimolar primer mixes of comaA-244f (a-f) and comaA-659r (a-f), and comaB-244f (a-f) and comaB-659r (a-f), respectively [7] (Table S2). Primers for each of comammox *Nitrospira* clades A and B *amoA* genes were initially tested in end-point PCR to check for a single dominant band in all amplifications, but subsequent qPCR was performed with clade A primers only because clade B primers produced no amplicons (data not shown). All qPCR amplifications were carried out as technical duplicates on a CFX96 Real-Time PCR Detection System (Bio-Rad, Hercules, CA, USA). Additional thermal cycling conditions and standards are described in the supplemental methods.

### 16S rRNA gene amplicon sequencing

The V4-V5 regions of 16S rRNA genes were amplified using primers 515F [37] and 926R [38] with Illumina adapters. The sequence data were produced by the US Department of Energy Joint Genome Institute (JGI) using standard operating procedures. Triplicates were combined, quantified, and sample amplicons were then pooled equally. The pooled library was sequenced on a MiSeq (Illumina) with 2×300 base reads. Sequence analysis was performed with QIIME2 version 2019.1.0 [39]. Sequences were trimmed to remove primer and adapter sequences with cutadapt [40]. Quality trimming, denoising, error-correction, paired-end read merging, chimera removal, and dereplication was performed with DADA2 [41], producing an amplicon sequence variant (ASV) table with 80,180-125,219 assembled sequences per sample (Supplemental file 1). The ASVs were taxonomically classified according to the SILVA database release 132 [42] using the scikit-learn classifier [43].

### Metagenome sequencing and analysis

Shearing of DNA, library preparation, and sequencing were performed at The Centre for Applied Genomics (TCAG) in Toronto, Ontario. Extracted DNA was quantified using the Qubit dsDNA HS Assay kit (Thermo Scientific, Waltham, MA, USA), and 500 ng of input DNA was sheared into ∼550 bp fragments using a LE220 Focused-ultrasonicator (Covaris, Woburn, MA, USA). Library preparation was performed using the TruSeq PCR-free Library Prep Kit (Illumina). Paired-end sequencing (2×250 bases) was performed on a HiSeq 2500 (Illumina) using the HiSeq Rapid SBS Kit v2 (500 cycle; Illumina), resulting in a total of ∼250 million paired-end reads with an average of 18.1 million paired-end reads per sample (Table S3).

Quality trimming and removal of adapter sequences were performed using AdapterRemoval version 2.2.2 [44] and the quality of the reads was checked with FastQC version 0.11.5 [45]. Reports were combined using MultiQC version 1.0 [46]. Open reading frames (ORFs) were predicted on the unmerged and unassembled trimmed forward reads using FragGeneScan-Plus [47]. Profile hidden Markov models (HMMs) for taxonomic marker (i.e., *rpoB*) and functional genes (i.e., *amoA*_AOA, *amoA*_AOB, and *nxrB*), downloaded from FunGene [48], were used to quantify the relative abundances and taxonomic affiliations of nitrifiers from the unassembled reads using MetAnnotate [49]. The HMM e-value threshold used by MetAnnotate for gene detection was 10^-3^, and the e-value threshold used for the USEARCH-based taxonomic classification step was 10^-6^. Within MetAnnotate, the database used for taxonomic classification was RefSeq release 80 (March 2017). MetAnnotate results were analyzed using the custom R script *metannotate_barplots.R* version 0.9 (available at https://github.com/jmtsuji/metannotate-analysis). To allow for approximate between-sample and between-HMM comparisons, hits for each HMM were normalized both to HMM length and total number of length-normalized HMM hits for *rpoB* (for more details, see the Github README).

### Assembly and binning of metagenome sequence reads

Sequence reads were processed through the ATLAS pipeline (version 2.0.6), which includes quality control, assembly, annotation, binning, bin dereplication, and read mapping [50, 51]; for further details, see supplemental methods. Taxonomy was assigned to bins according to the Genome Taxonomy Database [52] release 86, version 3, using the Genome Tree Database Toolkit version 0.2.2 [53]. Relative abundances of genome bins in metagenome sequence data were approximated by calculating the number of mapped reads to genome bins, normalized to the total number of assembled reads per metagenome. Each read required a minimum alignment score ratio of 0.908 (i.e., ∼95% sequence identity to the aligned region) to be considered and was mapped to up to 10 best aligned sites across the full genome set. Principal coordinate analysis (PCoA) ordination of samples used Bray-Curtis dissimilarities of relative abundance data using a custom R script.

### Analysis of genome bins

Average nucleotide identity (ANI) was calculated for the dereplicated AOA bin and *Nitrospira* bins compared to reference genomes using FastANI version 1.1 [54]. The *Nitrospira* bins were further searched for functional genes using reciprocal BLASTp [55] using the BackBLAST pipeline [56], version 2.0.0-alpha2 (doi:10.5281/zenodo.3465955). A concatenated core protein phylogeny, using a set of 74 core bacterial proteins, was generated to assess the phylogenetic placement of *Nitrospira* genome bins. The phylogenetic tree was constructed using IQ-TREE version 1.6.9 [57]. All predicted protein sequences from assembled contigs were compiled and clustered into 99% identity threshold groups within ATLAS; an HMM was used to identify AmoA sequences. *Nitrospira* AmoA were aligned along with reference sequences using MUSCLE [58] and a phylogenetic tree was built using MEGA7 [58]. Cyanase protein sequences (CynS) identified via BackBLAST were further analyzed phylogenetically as described further in the supplemental methods.

### Data availability

Amplicon sequence data are available on the JGI genome portal under sequencing project ID 1137811. Metagenome sequencing data and high completeness, low contamination genome bins classified to the *Nitrospirota* phylum were deposited in the European Nucleotide Archive under study accession number PRJEB30654.

## Results and Discussion

Prior to sequence-based analysis of RBC nitrifying biofilm sample communities, ammonia oxidizer cohort sizes were assessed using quantitative PCR (qPCR; Figure 2). The results showed distinct nitrifying community compositions, both along the treatment train (spatially; RBC 1 versus RBC 8) and between sampling time points (temporally; 2010 and 2016). Within each RBC train, the measured relative abundance of comammox *Nitrospira* genes, as well as the proportion of comammox *Nitrospira* within the total community, were consistently higher in RBC 1 than RBC 8 (Figure 2, Table S5), as were ammonium concentrations in water samples collected from below the sampled RBCs (Figure 1C, Table S4). Comammox *Nitrospira amoA* genes were more abundant than those of other ammonia oxidizers for most samples and time points (Figures 2, Figure S1, Table S5). In addition to comammox *Nitrospira*, the qPCR data also indicated a higher relative abundance of thaumarchaeotal 16S rRNA genes (inferred to represent AOA *Thaumarchaeota* based on other data) in RBC 8 than RBC 1 in the 2010 samples. In contrast, in 2016, thaumarchaeotal 16S rRNA gene abundances were higher in RBC 1 than RBC 8 for all four trains and were roughly equivalent in relative abundance to comammox *Nitrospira* (Figure 2). Although the abundances of AOB *amoA* genes were higher in RBC 1 than RBC 8 in both sampling years, the abundances of these genes were one to two orders of magnitude lower in 2016 than in 2010. Higher resolution sampling of all 2016 RBC stages showed the general trend of a decrease in *amoA* gene abundances across all four RBC flowpaths (Figure S1).

**Figure 2.**
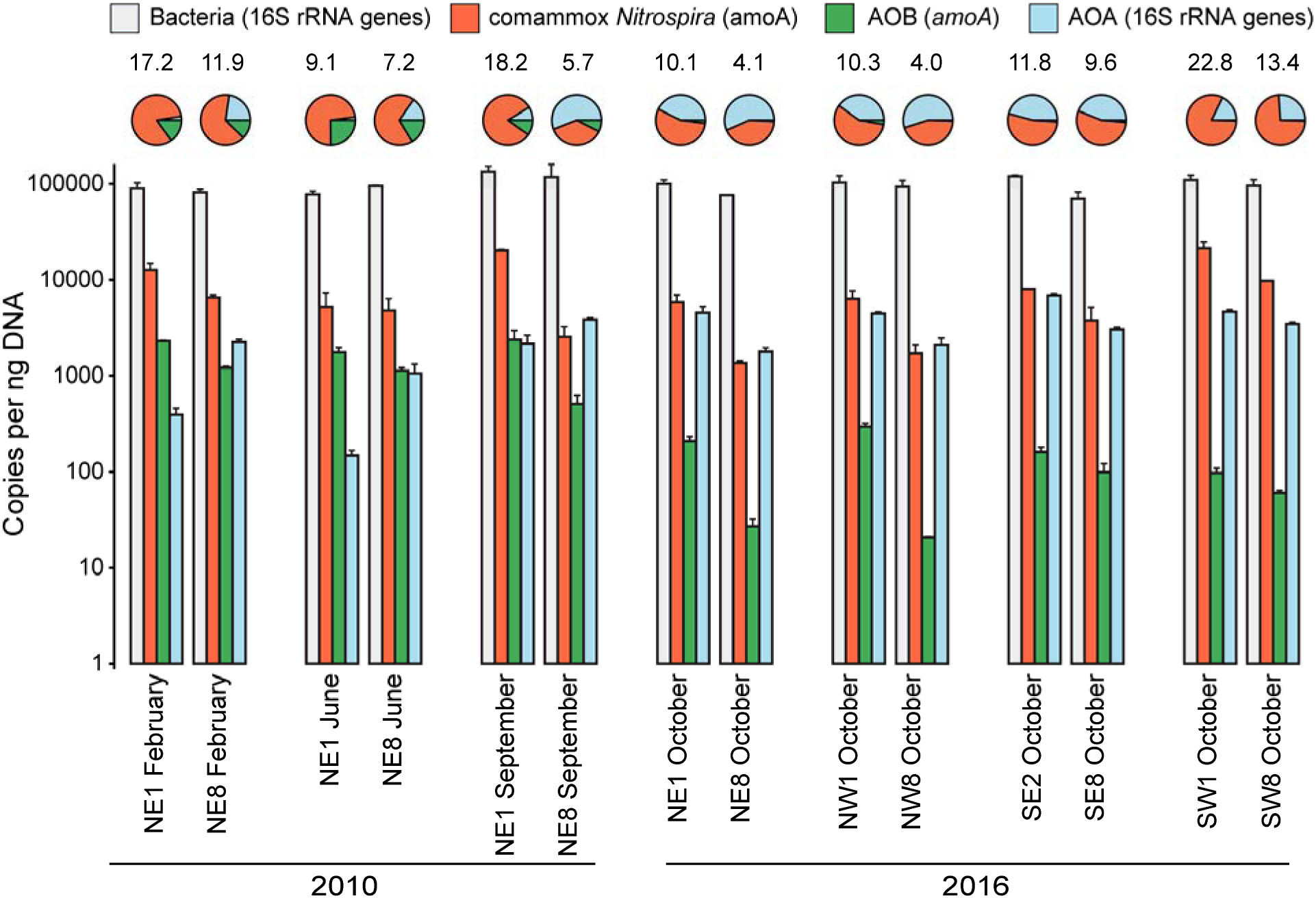
Bacterial 16S rRNA, comammox *Nitrospira* (*amoA*), AOB (*amoA*), and thaumarchaeotal 16S rRNA gene abundances for samples paired with metagenome sequencing. Error bars indicate standard deviation of technical qPCR duplicates. Pie charts show gene abundances as a proportion of all ammonia oxidizing prokaryotes. Numbers above the pie charts indicate the proportion (%) of the total community that all ammonia oxidizing prokaryotes represent. Gene copies were calculated based on the amount of DNA present in the original extractions but were not standardized to account for expected gene copy number per genome. NE, northeast train; NW, northwest train; SE, southeast train; SW, southwest train.

Although 16S rRNA genes cannot be used to differentiate between comammox *Nitrospira* and strict nitrite oxidizers [7], the overall relative abundance patterns of *Nitrospira* amplicon sequence variants (ASVs; Figure 3) were consistent with observations from qPCR data (Figure 2), showing higher relative abundances of *Nitrospira* in RBC 1 compared to RBC 8 for each sample pair. Eight ASVs present at ≥1% relative abundance were classified as *Nitrospira* (Figure 3, Supplemental file 1) and, taken together, these ASVs dominated all other populations detected in the RBC biofilm samples. In addition to *Nitrospira*, five ASVs present at ≥1% relative abundance were classified into the *Nitrosomonadaceae* family (Figure 3), and four corresponding to the genus *Nitrosomonas* affiliated only with the 2010 samples. The fifth *Nitrosomonadaceae* ASV affiliated with 2016 samples (ASV ID 4), but subsequent metagenome analysis suggested that this was not an ammonia oxidizer due to lack of a detectable *amo* gene system in the corresponding genome bin and phylogenetic distance from canonical AOB (data not shown). Thus, qPCR data were consistent with ASV data, with fewer AOB detected in 2016 samples overall. We detected a single thaumarchaeotal ASV that affiliated with the known AOA population within the RBCs (*Nitrosocosmicus* genus; [30]), and this ASV was present in all samples (Supplemental file 1) but at highest relative abundance (i.e., >1%) for 2016 samples (Figure 3), as expected based on qPCR data (Figure 2).

**Figure 3.**
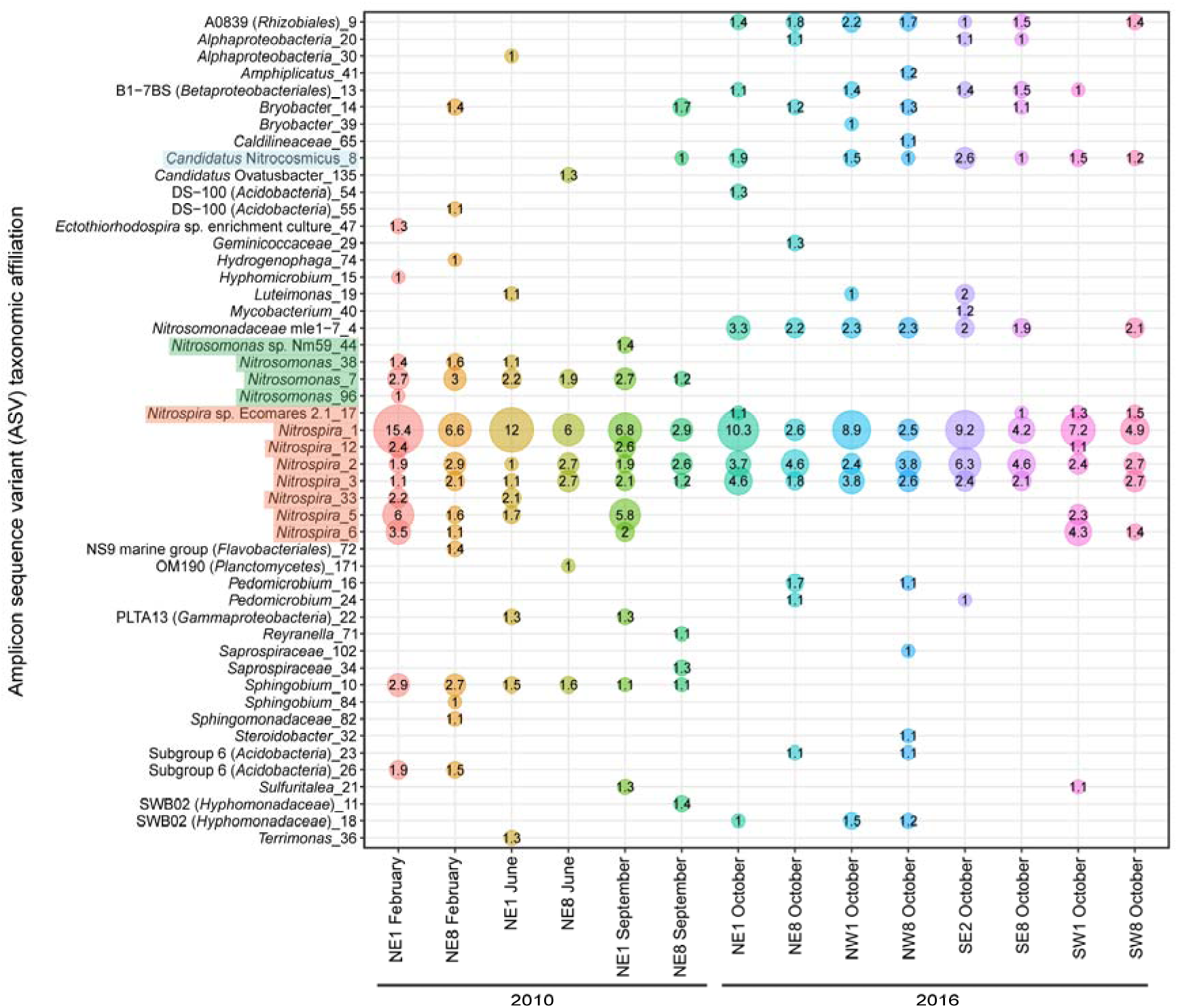
Relative abundances of amplicon sequence variants (ASVs) based on 16S rRNA gene amplicon sequencing, for samples paired with metagenome sequencing. Only taxa detected at ≥ 1% relative abundance are shown. The lowest level of informative taxonomic rank is given. Numbers after the underscore are the ASV number. The numbers inside the circles represent the relative abundance, in percent. *Nitrospira* ASVs are highlighted in orange, AOB ASVs are highlighted in green, and the AOA ASV is highlighted in blue.

Single copy taxonomic marker gene analysis of unassembled metagenome sequence data was consistent with qPCR and 16S rRNA gene sequence data, revealing that *Nitrospira* sequences dominated all samples and comprised between 8 and 32% of the total *rpoB* gene sequences among RBC samples from the two sampling years (Figure 4). The *Nitrospira*-affiliated *rpoB* gene relative abundances were higher in RBC 1 than RBC 8 for each train sampled. In addition, the normalized relative abundances of comammox *Nitrospira amoA* genes correlated well with the relative abundances calculated from qPCR data for the proportion of their *amoA* genes within the RBC biofilm communities (Spearman’s rank correlation, *r_s_* = 0.79, *p* < 0.001).

**Figure 4.**
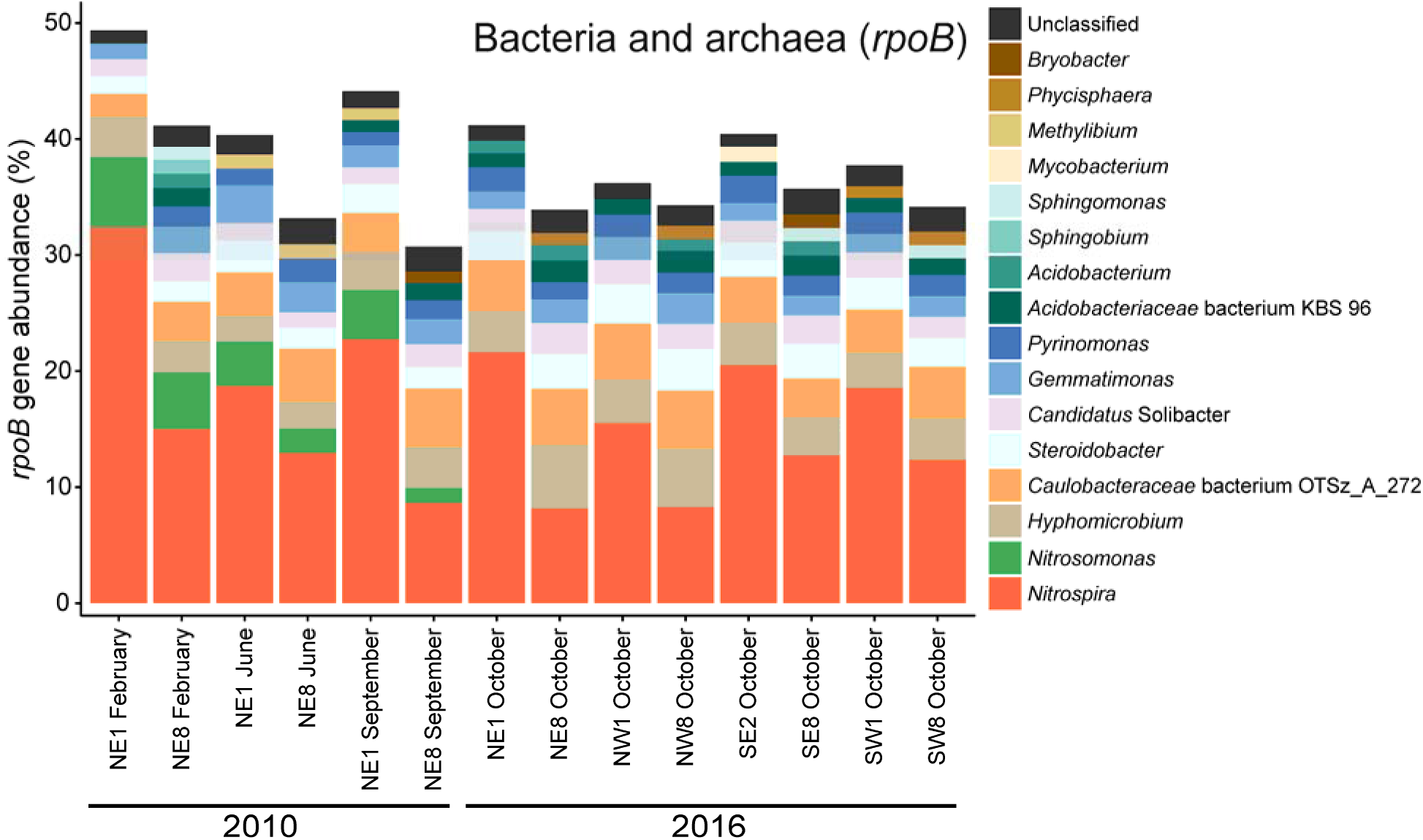
Taxonomic profiling of rotating biological contactor (RBC) microbial community by the hidden Markov model (HMM) for the RNA polymerase beta subunit (*rpoB*). Stacked bars represent the relative abundance of unassembled *rpoB* metagenome sequence reads classified at the genus level. Genera at greater than or equal to 1% relative abundance are shown. Dark orange and green shaded bars represent nitrifiers. *Nitrospira* species include both strict nitrite-oxidizing and comammox *Nitrospira*, because they cannot be distinguished at the genus level. NE, northeast train; NW, northwest train; SE, southeast train; SW, southwest train.

Normalized relative abundances of functional genes detected in unassembled read data allowed us to further estimate the relative community contributions of nitrifying community members. Assuming one *amoA* gene copy per genome, comammox *Nitrospira* represented between ∼5-30% of the total microbial community for all samples based on normalized *amoA* HMM hits (Figure 5B). *Nitrospira*-associated *rpoB* and normalized *amoA* gene abundances were similar (Figures 4, 5B), suggesting that the majority of *Nitrospira* present in the RBCs were comammox bacteria. The HMM search for *nxrB/narH* revealed that sequences affiliated with this model were also prevalent in RBC biofilm metagenomes, with combined hits at over 50% abundance normalized to total *rpoB* hits (Figure 5C). Based on phylogenetic placement of assembled genes in our metagenome sequence data that were detected using the *nxrB/narH* HMM, the majority of non-*Nitrospira* HMM hits likely represent denitrifying organisms encoding *narH* (data not shown), whereas *Nitrospira* HMM hits likely represent *nxrB*-encoding nitrifiers. The closest BLASTP search result for some non-*Nitrospira* affiliated *nxrB/narH* HMM hits was to a related protein (e.g., dimethyl sulfoxide reductase; data not shown), meaning that the normalized relative abundance we detected may be an overestimate of the true community contribution of *nxrB/narH*. Nevertheless, the high relative abundance of *Nitrospira*-affiliated *nxrB* gene sequences, which were more abundant in RBC 1 than RBC 8 metagenome sequences across all samples (Figure 5C), is further evidence for the dominance of comammox or NOB *Nitrospira* in the RBCs. Normalized relative abundances of *Nitrospira*-affiliated *nxrB* sequences that exceed the corresponding relative abundances of *Nitrospira*-affiliated *rpoB* sequences may be due to multiple copies of *nxrB* common to *Nitrospira* genomes [59], in contrast to single copies common for *rpoB* genes [60], or may be due to known biases that occur during HMM hit count normalization.

**Figure 5.**
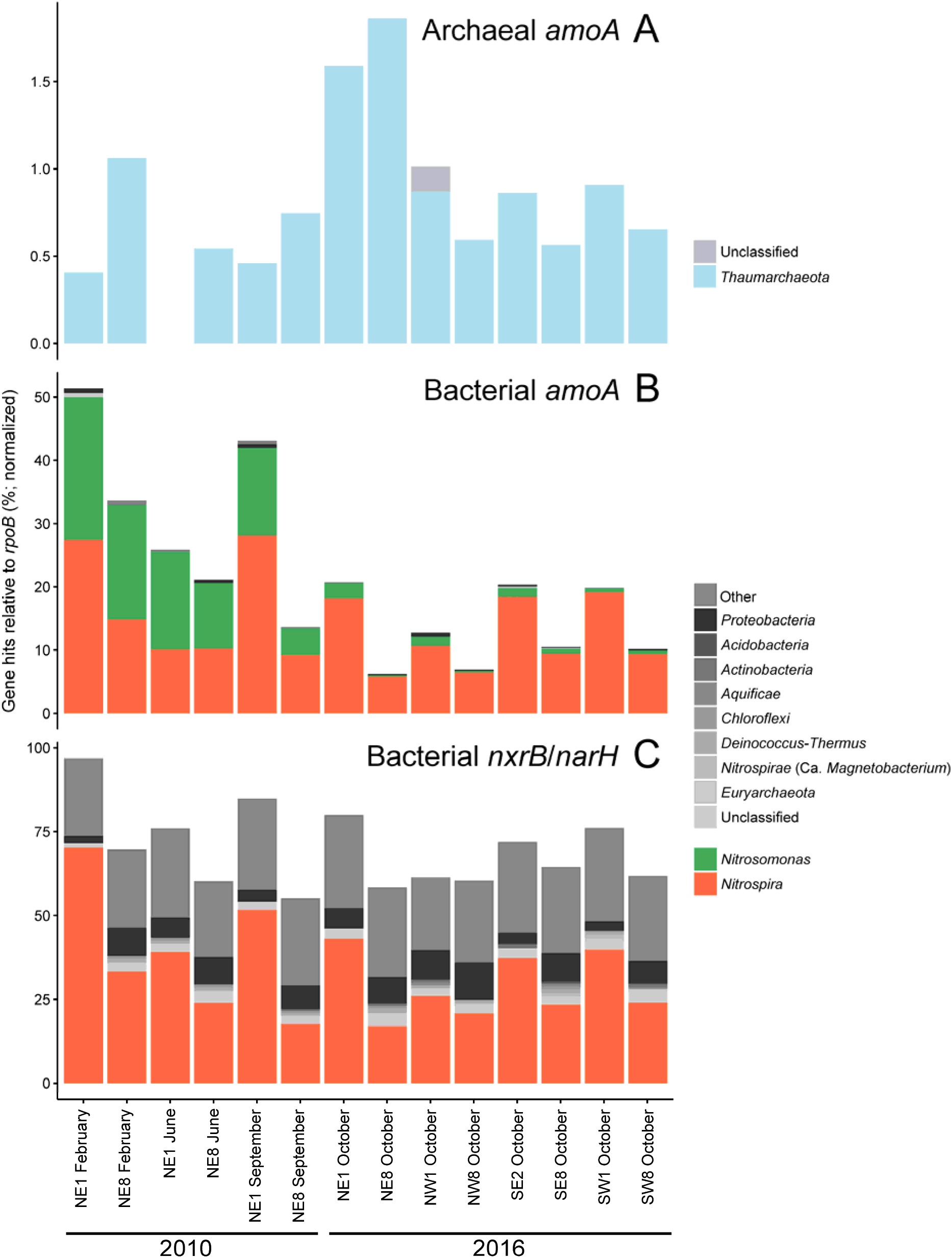
Functional gene profiling of nitrifiers in rotating biological contactor (RBC) microbial communities by hidden Markov models (HMMs). Stacked bars represent the normalized abundances of functional genes, relative to *rpoB*, from unassembled metagenome sequence reads. Genera at ≥1% normalized relative abundance are shown with taxa labels, along with genera at <1%, which are marked as “Other” at the uppermost grey bar. Blue, green, and orange bars represent nitrifiers. Other genera at ≥1% normalized relative abundance are collapsed to the phylum level for clarity and are shown in grey. (A) HMM search for the *amoA* gene of ammonia-oxidizing archaea, which are grouped at the phylum level (*Thaumarchaeota*) for clarity. No HMM hits were found for sample NE1 June 2010. (B) HMM search for the *amoA* gene of ammonia-oxidizing bacteria and comammox *Nitrospira*. (C) HMM search for the *nxrB* gene of *Nitrospira* and other NOB. This HMM also may have detected genes for nitrate reductase (*narH*). NE, northeast train; NW, northwest train; SE, southeast train; SW, southwest train; *amoA*, ammonia monooxygenase subunit A gene; *nxrB*, nitrite oxidoreductase; *narH*, nitrate reductase.

Consistent with qPCR data, *Nitrosomonas*-affiliated *rpoB* gene sequences were detected at ≥1% in the 2010 samples only. The *rpoB* gene of AOA was not detected at ≥1% relative abundance in any of the samples but ranged from 0.01-0.3% across all samples (data not shown). The HMM search for the *amoA* gene of AOA similarly revealed low relative abundance in the metagenome dataset (<2% relative to *rpoB* for all samples; Figure 5A). For 2010 samples, *amoA* gene sequences affiliated with *Thaumarchaeota* were more abundant in RBC 8 than RBC 1 samples, whereas in 2016 they were more abundant in RBC 1 than RBC 8 samples, except in the NE 2016 sample. The HMM hits for AOA *amoA* genes in the unassembled metagenome sequences were low, with only 64 hits to this HMM for all samples combined, but the AOA *amoA* normalized relative abundance pattern was nevertheless correlated with the corresponding proportion of AOA 16S rRNA genes within the total community determined via qPCR (Spearman’s rank correlation, *r_s_* = 0.67, *p* = 0.009). The *amoA* HMM for AOB detected sequences of both comammox *Nitrospira* and AOB. Overall, the relative abundances of AOB *amoA* genes were higher in RBC 1 than RBC 8 of the corresponding sample pairs, in both sampling years (Figure 5B). In 2010, *Nitrosomonas* spp. and *Nitrospira* spp. were at roughly equal relative abundances, whereas *Nitrospira* spp. were the dominant ammonia oxidizers detected in 2016 metagenome sequences. The *amoA* genes of the AOB *Nitrosospira* were also detected, but always at <1% relative abundance. Overall, the relative abundance patterns for AOB *amoA* genes correlated well with their corresponding relative abundances determined by qPCR for the proportion of their *amoA* genes within the total community (Spearman’s rank correlation, *r_s_* = 0.97, *p* < 0.001).

Metagenome-assembled genomes (MAGs) obtained from RBC metagenome sequence data were also examined for the temporal and spatial distributions of *Nitrospira* populations. Summed together, *Nitrospira* MAGs (see taxonomic classification description below) had a higher relative abundance in RBC 1 than RBC 8 for each sample pair (Figure 6). This pattern was also the case for the bins classified as clade A comammox *Nitrospira*, which comprised a majority of recruited reads to *Nitrospira* MAGs across most samples. At the same time, individual *Nitrospira* MAGs displayed distinct temporal and spatial distributions. Several of the comammox MAGs had different abundance patterns in 2010 than in 2016. Specifically, RBC035, RBC100, and RBC069 were present at higher relative abundances in 2010 samples than in 2016 samples, and relative abundances of RBC001 and RBC083 were generally higher in 2016 than 2010 samples (Figure 6, Supplementary file 2). Multiple comammox *Nitrospira* populations were found in the same RBC, and this high diversity of comammox *Nitrospira* contrasts with the observed low AOA diversity (i.e., only one *Nitrosocosmicus* species; see below) for all RBCs sampled. Although 0.5-3.1% (median 1.1%) of assembled reads associated with *Nitrospira* MAGs were mapped to more than one MAG (data not shown), this level of crossover read mapping is insufficient to explain the high observed diversity of comammox *Nitrospira* for most RBCs, meaning that the high observed diversity is likely not an artefact of read mapping. Using all of the dereplicated bins, the overall microbial communities of the samples group by year and RBC number (Figure S2B), similar to an ordination prepared from 16S rRNA gene sequence data (Figure S2A). This demonstrates that distinct microbial communities were present in each of the two sampling years and along the RBC flowpaths.

**Figure 6.**
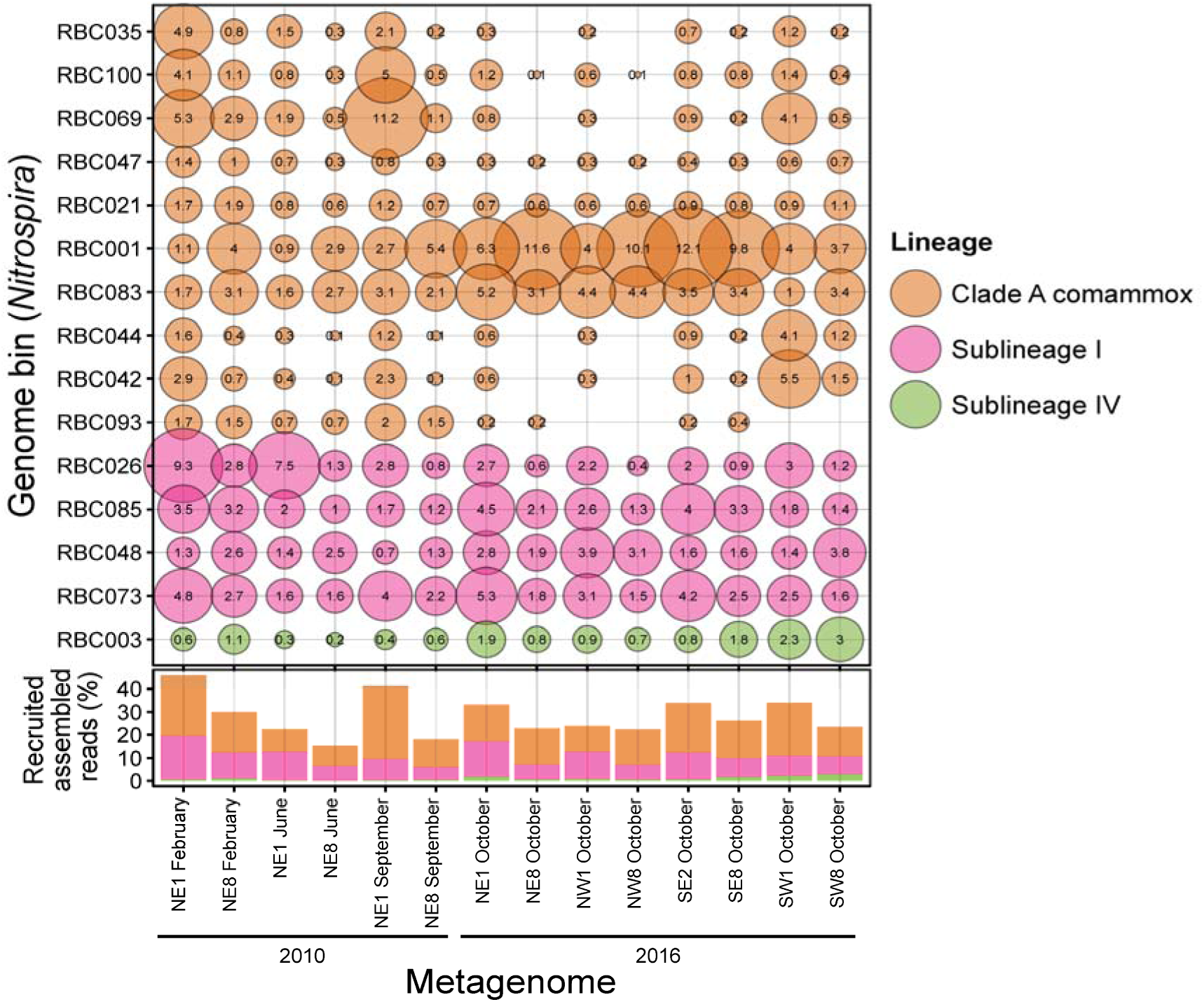
Relative abundance of *Nitrospira* bins across samples based on total recruited reads. The size of the bubbles indicates the proportion (%) of assembled reads from each sample that mapped to each genome bin. Stacked bars summarize the total proportions of assembled metagenomic reads recruited to the genome bins.

The dominance of comammox *Nitrospira* in biofilm samples collected from RBCs treating municipal wastewater of Guelph, Ontario, Canada is in contrast to several previous studies of full-scale activated sludge WWTPs that report comammox *Nitrospira* at lower relative abundances than AOA and other AOB [7, 18–21], albeit for systems with ammonium concentrations and operational parameters distinct from those in the RBCs. This current study, along with other recent studies [12, 15, 17, 18, 22], shows that comammox *Nitrospira* can dominate municipal wastewater treatment systems. Although previous analyses of the Guelph WWTP did not test for these newly discovered ammonia oxidizers [29, 30], the presence of comammox *Nitrospira* was predicted previously from activity data and an abundance of *Nitrospira* cells that actively assimilated labelled bicarbonate when incubated with ammonium [30]. The current discovery of abundant comammox *Nitrospira* communities in Guelph RBCs suggests a correspondingly important role for comammox within the 440,000 m^2^ of RBC biofilm associated with this full-scale wastewater treatment system.

High relative abundances of comammox *Nitrospira* have been reported for other engineered environments with relatively low ammonium levels, such as drinking water treatment systems. For example, comammox *Nitrospira* were found in a drinking water treatment plant [8] and distribution systems [10], and they were more abundant than other ammonia oxidizers in a groundwater well studied previously [7]. Associated with drinking water treatment plants, comammox *Nitrospira* were the most abundant nitrifiers in groundwater fed rapid sand filters [13]. In those filters, clade B comammox *Nitrospira* dominated, in contrast to clade A members that we detected in the sampled RBCs. A study examining recirculating aquaculture system biofilters found that comammox *Nitrospira* were more abundant than AOA and AOB [61]. Along with these other studies, the dominance of comammox *Nitrospira* in the RBCs suggests that they compete well in engineered environments with relatively low ammonium concentrations. At the Guelph WWTP, the RBCs are located downstream of activated sludge aeration basins. Ammonium concentrations entering this tertiary treatment system are approximately two orders of magnitude lower than within aeration basins. For example, on the day of sampling in 2016, the ammonium concentration in aeration basin influents averaged 2.5 mM NH_3_-N, which is typical of aerobic secondary aeration basin conditions reported elsewhere [e.g. 20–22, 62]. A large surface area for attached growth may be an important factor explaining the dominance of comammox bacteria, given their high prevalence in wastewater treatment systems with attached growth components [18]. The predicted low ammonium niche and biofilm-specific growth of comammox *Nitrospira* [4, 5] may help explain the dominance of these nitrifiers in the tertiary treatment system RBCs in Guelph, Ontario.

Although relatively low ammonium concentrations likely contribute to the overall success of comammox *Nitrospira* within the RBC biofilm samples, a gradient of decreasing ammonium along the RBC flowpaths could also account for relative abundance changes of nitrifying microbial communities, both temporally and spatially. We predicted initially that comammox *Nitrospira* abundances in RBC 8 samples would exceed those in RBC 1 samples due to the predicted low ammonium niche of comammox *Nitrospira* [4], but the opposite trend was observed for both sample years (Figures 2, 5, 6). However, ammonium concentrations below RBC 1 were already relatively low (e.g., <18 µM for all 2016 samples; Figure 1C), and it may be that ammonium concentrations in RBC 8 influent (e.g., <4 µM for all 2016 samples; Figure 1C) were below the optimum for comammox *Nitrospira*. Consistent with an “oligotrophic lifestyle”, Kits *et al*. [63] reported a high apparent affinity of *Ca.* Nitrospira inopinata for ammonium (K_m(app)_ of 0.65 to 1.1 µM total ammonium), and V_max_ reached by ammonium concentrations of ∼5 µM, which would be similar to ammonium concentrations observed for water samples collected from RBCs sampled in this study, especially for RBC 1 stages. The affinities for ammonium of the comammox *Nitrospira* detected in the RBCs are unknown but may be similar those reported for *Ca.* N. inopinata.

Ammonium concentrations alone may not explain comammox *Nitrospira* abundances, however. In contrast to published studies reporting low abundances of comammox *Nitrospira* in activated sludge flocs [7, 19–21], a recent PCR-based study found that comammox *Nitrospira amoA* genes were more abundant than those of AOB in activated sludge samples from eight separate WWTPs, despite relatively high ammonium concentrations [22]. This suggests that, as seen for AOA, [e.g. 64–68], the abundances of comammox *Nitrospira* vary among WWTPs. More studies are needed to identify factors that affect comammox *Nitrospira* abundances, such as substrate range, oxygen requirements, growth rates, growth yields, and biofilm formation capabilities [69].

In addition to targeting nitrification-associated genes to identify comammox *Nitrospira* in extracted DNA or unassembled metagenome sequences, we explored *Nitrospira* diversity within assembled and binned metagenome sequence data. From a final dereplicated dataset with 101 genome bins (Supplementary file 2), 15 MAGs were classified within the *Nitrospirota* phylum. These MAGs all had high completeness (≥75%) and low contamination (<5%; Table S6). All but one of these *Nitrospirota* MAGs were classified under the Genome Taxonomy Database (GTDB) taxonomy within the *Nitrospira* or *Nitrospira* A genera of the *Nitrospiraceae* family; the final MAG (*Nitrospira* bin RBC003) was classified within the uncultured UBA8639 lineage (genus and family name) of the *Nitrospirales* order. Using a concatenated set of 74 core bacterial proteins, these MAGs were further classified according to traditional *Nitrospira* phylogeny (Figures 7, S3). A majority (10 of 15) of the *Nitrospirota* MAGs clustered within clade A comammox *Nitrospira*, four clustered within sublineage I (strict nitrite-oxidizing bacteria), and one (which was classified to the UBA8639 lineage according to GTDB taxonomy) clustered within sublineage IV (strict nitrite-oxidizing bacteria) of the *Nitrospira*. Thus, the set of 15 *Nitrospirota* MAGs will be referred to as *Nitrospira* MAGs for conciseness. This high diversity of *Nitrospira* MAGs is consistent with multiple *Nitrospira* ASVs detected with 16S rRNA gene amplicon sequencing (Figure 3). Using average nucleotide identity (ANI) analysis, the 15 *Nitrospira* MAGs were compared to one another and to *Nitrospira* genomes from enrichment cultures and metagenomic surveys (Figure S3). Only one MAG, RBC035, had ≥95% ANI to previously recovered comammox genomes (*Nitrospira* bin UW_LDO_01 [27] and *Candidatus* Nitrospira nitrosa [6]); all other comammox *Nitrospira* MAGs had <95% ANI to other reference genomes. This suggests that these MAGs represent distinct comammox *Nitrospira* species from most other previously described comammox species.

**Figure 7.**
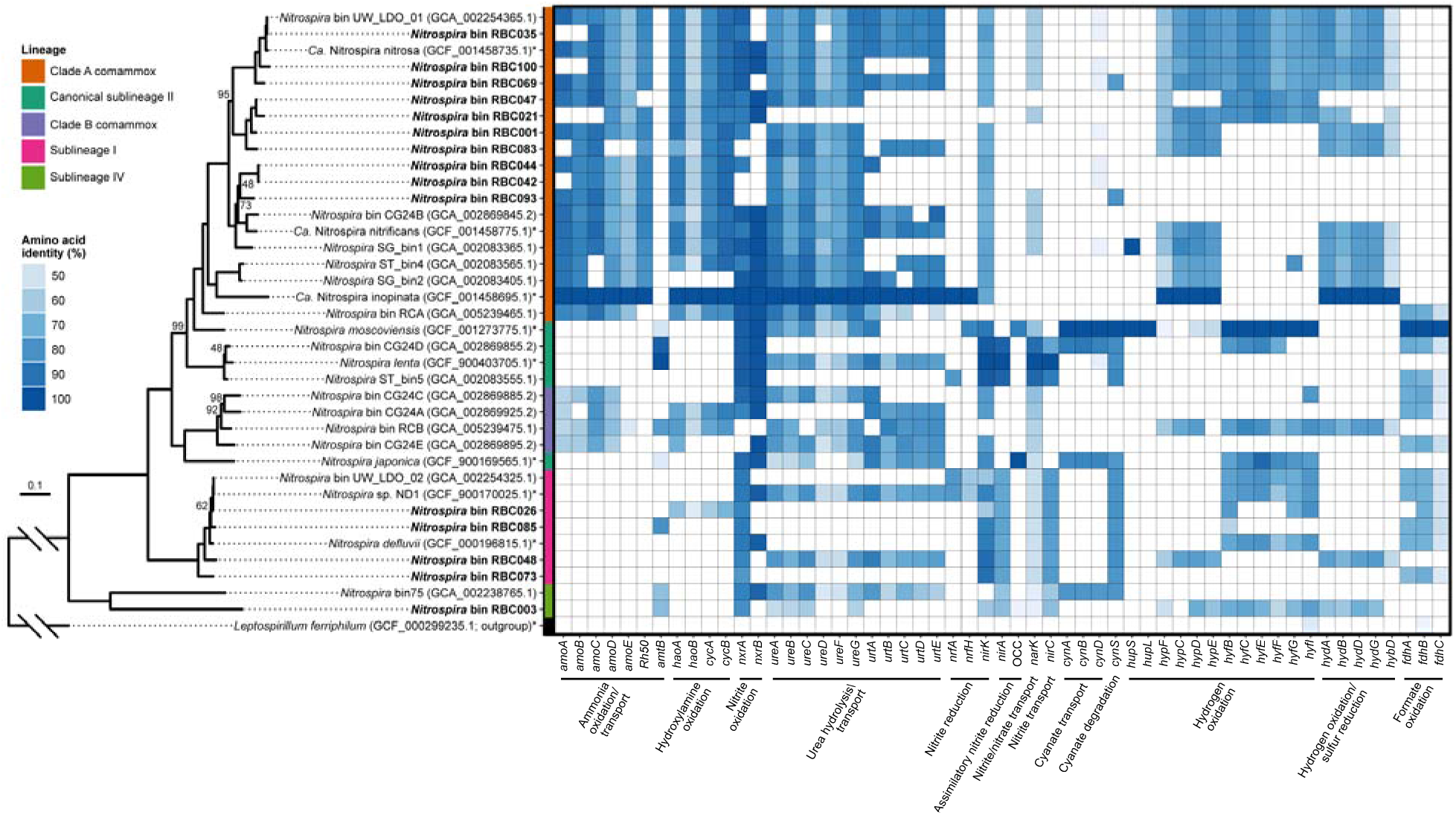
Phylogeny and gene pathways in *Nitrospira* bins and reference genomes. The bins recovered from this study are indicated in bold, an asterisk indicates that the genome is from an enrichment or pure culture. Phylogenomic analysis was performed using a concatenated set of 74 core bacterial proteins. All bootstrap values are 100% except those shown, and the scale bar represents the proportion of amino acid change. Gene annotation was determined using reciprocal BLASTP against four reference *Nitrospira* genomes (*Ca.* N. inopinata, *Nitrospira moscoviensis*, *Nitrospira lenta, Nitrospira japonica*), and amino acid identity to the reference gene is indicated in the heatmap. Abbreviations: *amo*, ammonia monooxygenase; *Rh50*, ammonia transporter; *amt*, ammonia transporter; *hao,* hydroxylamine oxidoreductase; *cyc*, cytochrome c; *nxr*, nitrite oxidoreductase; *ure*, urease; *urt*, urea transporter; *nrf,* cytochrome c nitrite reductase; *nirK*, dissimilatory nitrite reductase; *nirA*, assimilatory nitrite reductase; OCC, octaheme cytochrome c; *narK*, nitrite/nitrate transporter; *nirC*, nitrite transporter; *cynABD,* cyanate transporter; *cynS*, cyanate hydratase; *hup*, group 2a [Ni-Fe] hydrogenase; *hyp*, hydrogenase accessory protein; *hyf,* putative group 4 hydrogenase; *hyb* and *hyd*, group 3 [Ni-Fe] sulfur-reducing hydrogenase; *fdh*, formate dehydrogenase.

Based on 99% identity clustering of amino acid sequences, all assembled *amoA* genes of *Nitrospira* spp. grouped phylogenetically within comammox *Nitrospira* clade A (Figure S4), aside from RBC group F (a truncated sequence which was dropped from phylogenetic analysis). This is consistent with other studies reporting comammox *Nitrospira* in WWTPs [7, 12, 14–18]. Clade B comammox *Nitrospira* were not detected and the factors affecting the presence and absence of clades A and B remain unknown. Relatedness of *Nitrospira* bins based on *amoA* phylogeny (Figure S4) was similar to that obtained with the concatenated protein set, but not identical (Figures 7, S3). Recently, Xia *et al.* [14] proposed that clade A could be further subdivided into clade A.1, which includes the cultivated comammox *Nitrospira*, and clade A.2, which contains only uncultivated comammox *amoA* sequences, including those from drinking water systems (OQW38018 and OQW37964) [10]. The *Nitrospira amoA* gene sequences detected in the RBCs were classified into both of these clade A subdivisions (Figure S4). Translated sequences of clade A.2 *amoA* genes from the RBCs were most closely related to *amoA* sequences obtained from metagenome sequences of drinking water systems [10] or to a clone library sequence from a rice paddy (Genbank accession AKD44274). Several comammox *Nitrospira* related to the rice paddy clone have been detected in other full-scale WWTPs, and this clade was described as a new clade A cluster by the authors [22].

Using a reciprocal BLASTP analysis, the *Nitrospira* MAGs were further evaluated for the presence of *amo, hao,* and *nxr* genes to assess potential contributions to nitrification [5, 6]. *Nitrospira* bin RBC047 contained all the genes necessary to perform complete ammonia oxidation, although it was missing an ammonium transporter gene (Figure 7). *Nitrospira* bins RBC069, RBC001, and RBC044 were missing *nxrB* genes but contained the other genes for ammonia and nitrite oxidation and had a Rh50 type ammonium transporter. In total, there were five clade A comammox MAGs that contained the *amoA* gene, which is used as a marker gene for comammox *Nitrospira*. The other five MAGs that clustered phylogenetically within clade A comammox *Nitrospira* contained other genes for ammonia oxidation, supporting their phylogenetic placement on the tree. The other *Nitrospira* MAGs contained genes only expected in NOB, except bin RBC026, which had genes for hydroxylamine oxidation. These genes had >90% amino acid identity to hydroxylamine oxidation genes from known *Nitrosomonas* spp., implying that the genes were present in bin RBC026 due to mis-binning of the associated ∼4.3 kb contig. Also, the high (93%) ANI between bin RBC026 and comammox bin RBC083 suggests that cross-contamination of these bins may be possible (Figure S3). Overall, one *Nitrospira* MAG contained the entire gene pathway required for comammox, four bins contained only genes expected for NOB, and nine bin clusters could potentially represent comammox *Nitrospira* but lacked the complete gene pathway. Cells of *Nitrospira* within RBC biofilm samples were previously demonstrated to assimilate labelled bicarbonate in the presence of ammonium [30], indicating autotrophic activity of these bacteria and suggesting the possibility of ammonia oxidation.

High functional diversity was observed in the RBC *Nitrospira* MAGs. In the current study, the presence of a nearly complete *ure* operon (Figure 7) indicates that comammox *Nitrospira* of the RBCs could hydrolyze urea to ammonia, which they could then use for ammonia oxidation. The presence of these genes is consistent with clade A comammox *Nitrospira* genomes and enrichment cultures that use urea [5, 6, 69, 70]. Because the comammox *Nitrospira* bins did not contain any *fdh* genes for formate dehydrogenase (Figure 7), these RBC populations likely cannot use formate as an alternative electron donor, which can be used by NOB *Nitrospira* and most clade B comammox bacteria [70–73]. Most clade A comammox bacteria described so far lack the genes for formate dehydrogenase [70], with the exception of *Nitrospira* bin RCA [74]. Also consistent with previous findings [18, 70], several of our comammox *Nitrospira* MAGs contained partial pathways for group 3b [Ni-Fe] sulfur-reducing hydrogenase genes (*hyb* and *hyd* genes; Figure 7), which indicates that hydrogen may serve as an alternative electron acceptor for some of these comammox *Nitrospira*. Although several of these MAGs also encoded genes for a putative group 4 hydrogenase (*hyf*), the amino acid sequences of the putative large subunit hydrogenase genes (*hyfG*) did not contain the two CxxC motifs needed for ligating the NiFe center, implying a potential alternative function of these genes other than hydrogen oxidation, as has been suggested previously for other bacteria [75].

Multiple clade A comammox *Nitrospira* MAGs contained genes for accessory proteins for hydrogenases (*hyp*) but none of the *Nitrospira* MAGs contained group 2a [Ni-Fe] hydrogenase genes (*hup*), as seen in the NOB *Nitrospira moscoviensis* [76]. All of the comammox *Nitrospira* MAGs lacked the gene for assimilatory nitrite reduction (*nirA*).

Surprisingly, two clade A comammox *Nitrospira* bins (RBC069 and RBC093) that contained genes for ammonia oxidation also encoded the cyanase gene (*cynS*; Figures 7, 8). Prior to this observation, only canonical NOB *Nitrospira* were known to possess this gene [70, 73, 74, 77, 78]. The *cynS* genes were on long contigs (i.e., 182 kb and 121 kb, RBC069 and RBC093, respectively), and the primary sequences of most genes adjacent to the *cynS* genes, when queried against RefSeq using BLASTP, were most similar to genes belonging to known comammox *Nitrospira*. This implies that the presence of *cynS* in RBC comammox MAGs was not caused by an error in genome binning. Both cyanase genes were distinct from those of other strict NOB *Nitrospira,* sharing <75% identity (blastn) to *Nitrospira* homologues in the non-redundant nucleotide database. Nevertheless, the predicted primary sequences of the *cynS* genes contained the key active site residues for cyanases (Arg96, Glu99 and Ser122) proposed previously [79], further supporting the functional role of the enzymes (Figure 8A).

**Figure 8.**
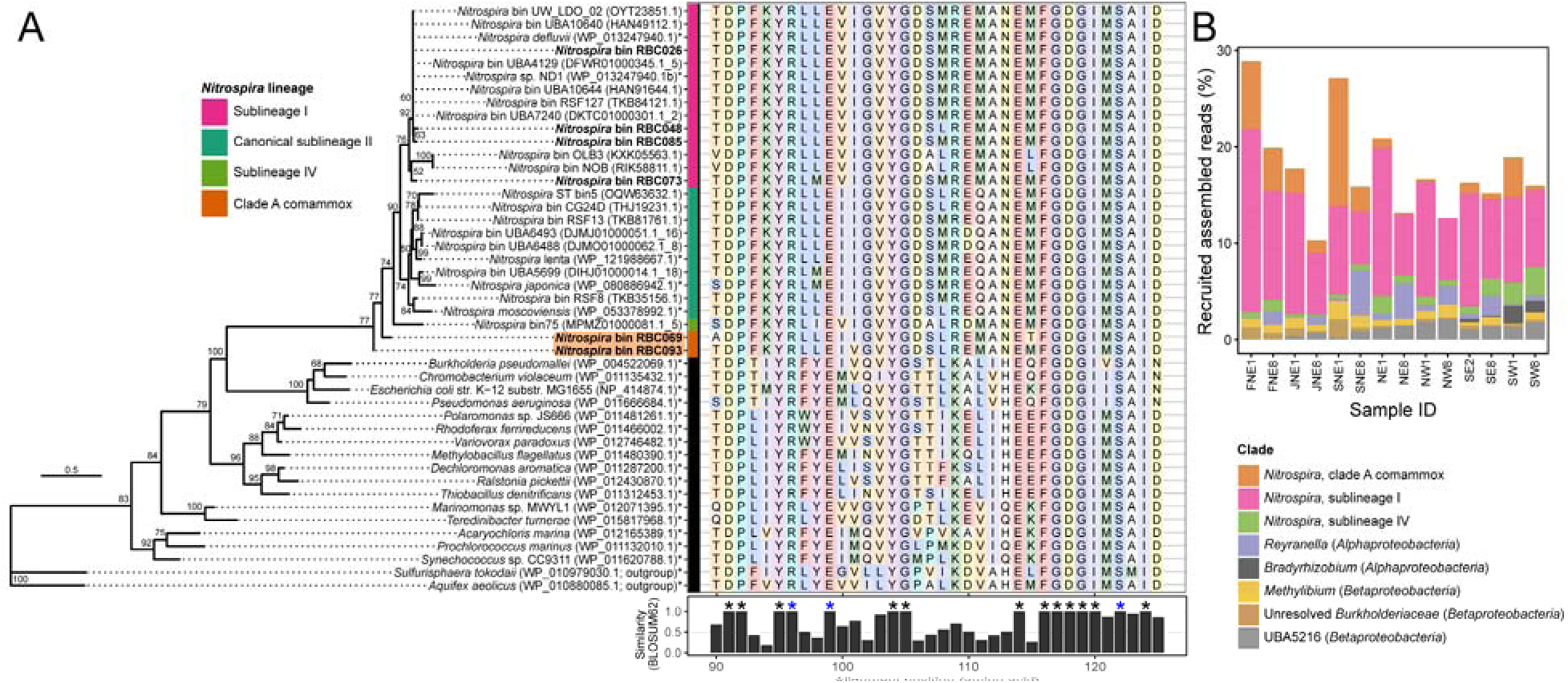
Phylogeny and relative abundances of cyanases (CynS) detected in RBC metagenome sequencing data. (A) Maximum likelihood phylogeny of CynS primary sequences detected in *Nitrospira* genome bins from this study compared to reference sequences from NCBI. Sublineages of the *Nitrospira* clade are highlighted by the coloured vertical bar, including potential clade A comammox bacteria, which are highlighted in orange. Genome bins recovered from the RBCs are bolded, and an asterisk appears after the names of genomes of cultivated organisms. Bootstrap values over 50% are shown, and the scale bar represents the proportion of amino acid change. To the right of the phylogeny, a portion of the CynS primary sequence alignment is shown along with overall sequence similarity based on the BLOSUM62 metric. Residues with 100% conservation are marked with an asterisk. Blue asterisks are used to mark known catalytic residues of CynS, namely Arg96, Glu99, and Ser122. (B) Relative abundances of all RBC genome bins containing *cynS* among RBC metagenomes. Genome bins are collapsed at the genus level into stacked bars, and relative abundances are based on the proportion (%) of mapped assembled reads to genome data.

Interestingly, the phylogenetic placement of the comammox cyanases was basal to all other known cyanases among canonical NOB *Nitrospira* (Figure 8A). This basal placement does not match housekeeping gene phylogenies of *Nitrospira* (e.g., Figure 7), indicating the potential for lateral gene transfer for cyanase acquisition. Indeed, we detected two toxin-antitoxin genes in the vicinity of the cyanase gene for RBC093, supporting the potential for a recent lateral gene transfer event [27]. Bins containing *cynS* together represent a substantial proportion of the RBC microbial community (i.e., ∼10-30%; Figure 8B). If these genes are expressed and produce functional gene products, our results indicate that *Nitrospira*, including novel *cynS*-containing comammox *Nitrospira*, dominate cyanate degradation to ammonia within RBC biofilm for a direct supply of ammonia (in the case of comammox *Nitrospira*) or for supplying ammonia to other nitrifiers through reciprocal feeding [77].

The comammox *Nitrospira* in the RBCs lack traditional cyanate transporters (*cynABD*), though one cyanate transporter gene (*cynD*) seemed to have moderate homology to another non-orthologous ABC transport gene, as indicated by the low amino acid identity hits to *cynD* among certain *Nitrospira* (Figure 7). These hits may represent another ABC transport protein, a nitrate/sulfonate/bicarbonate ABC transporter (accession number WP_121988621). If the comammox *Nitrospira* are unable to transport cyanate, our results may suggest that these *Nitrospira* degrade cyanate generated internally, for example from the degradation of carbamoyl phosphate or from a spontaneous isomeric conversion of urea to cyanate [80–84]. However, it is also possible that alternate transporters could be used for uptake of cyanate, such as nitrate permease [85], or the nitrite/nitrate transporter NarK, which is proposed to also transport cyanate [77]. If confirmed, this would represent an undescribed method of nitrogen uptake and utilization for comammox bacteria.

Multiple populations of comammox *Nitrospira* within the same RBC system suggests that these nitrifiers do not compete directly but instead occupy unique niches within the RBCs as a result of encoded functional diversity, for example because of distinct energy sources (e.g., hydrogen, urea, cyanate) or microenvironments within the biofilm that vary in oxygen or ammonium concentrations. Indeed, the abundances of sequences that recruited to comammox *Nitrospira* (based on read recruitment to bins) varied temporally and spatially among RBCs (Figure 6), implying that complex physicochemical factors govern comammox in the RBCs. Several RBC comammox *Nitrospira* MAGs may even represent closely related strains (Figure S3), adding further interest to understanding the factors influencing the distribution and ecology of these bacteria.

Genome bins of other ammonia oxidizers were also recovered. Following dereplication, four bins were classified as *Nitrosomonas* (Supplementary file 2). These populations were present at a higher relative abundance in 2010 samples than 2016 samples. A single dereplicated *Nitrosocosmicus* (AOA) bin was detected (RBC071; Supplementary file 2), which indicates the presence of a single *Thaumarchaeota* population and matches previous data from the RBCs [29, 30]. In 2010, this species was present at a higher relative abundance in RBC 8 than RBC 1, with the opposite pattern occurring in 2016 samples. Classified as *Ca.* N. exaquare under GTDB taxonomy, the AOA bin had an ANI of 96.1% between it and the *Ca.* N. exaquare genome (GenBank accession number CP017922.1), which indicated that this AOA MAG corresponded to the same archaeon enriched and characterized from an RBC biofilm sample.

The observed pattern of AOA *amoA* abundances for 2010 samples analyzed in this study (Figure 2, Figure 5A) was consistent with previous data obtained from separate DNA extractions of the same samples [29], demonstrating that AOA were indeed more abundant in RBC 8 than RBC 1 samples in 2010. Although AOA in 2016 samples were at a lower relative abundance in RBC 8 compared to RBC 1, this matches data from more recent samples collected from the RBCs in December 2015 [30]. Lower ammonium concentrations in recent years (Figure S5) may have shifted the *Ca*. N. exaquare AOA population toward RBC 1, and cleaning of the RBC trains in 2014 would have impacted the microbial communities. Lower ammonium concentrations beneath RBCs and RBC cleaning may have impacted AOB populations as well. Both qPCR (Figure 2) and metagenome sequence data (Figure 5B) indicated an overall decrease in AOB *amoA* gene relative abundances in the 2016 samples compared to 2010. Most AOB have a relatively low ammonia affinity [63, 86, 87] and may be outcompeted when ammonium concentrations are low by ammonia oxidizers with higher ammonia affinities (i.e., comammox *Nitrospira* and AOA).

### Conclusions

This study used complementary methods to demonstrate that comammox *Nitrospira* dominate microbial communities in biofilms from this engineered WWTP system. Detected comammox *Nitrospira* exhibited a high level of taxonomic and functional diversity, in contrast to the single population of AOA found in the same systems. The evolutionary history of comammox *Nitrospira* prior to establishing within the WWTP, perhaps within heterogeneous soil or sediment samples, may be a factor that contributed to the high diversity observed within RBCs. There is some phylogenetic evidence that the AOA selected within the RBCs may have originated on skin surfaces [30, 88], which would offer relatively low habitat heterogeneity. Additionally, other studies have found the coexistence of closely related *Nitrospira* in WWTPs [17, 89, 90], suggesting that *Nitrospira* may be able to occupy many different niches in engineered nitrifying environments. Although comammox *Nitrospira* were numerically dominant, their contributions to nitrification in the RBC environment remain unclear. Future studies using differential inhibitors, isotope tracer studies, and laboratory cultivation may be useful for elucidating the contributions of comammox *Nitrospira* to nitrification in the RBC biofilms. In addition, the comammox *Nitrospira* that fall within the newly proposed clade A.2 that lacks cultured representatives [14], along with the *cynS*-containing comammox populations, would be ideal targets for cultivation.

The Guelph WWTP RBCs represent a unique and useful system to study the ecology of comammox *Nitrospira* as a result of inbuilt environmental gradients and the combined presence of comammox *Nitrospira*, AOA, AOB, and NOB. Tertiary treatment systems such as these RBCs can be used to produce high quality effluents, which may be particularly important when WWTPs discharge into environments that have low assimilative capacities or are ecologically sensitive. A longer-term study of the microbial community and operating parameters (e.g., hydraulic retention time, oxygen concentration, organic loading) would help elucidate the factors that affect the microbial community, with a focus on the ammonia oxidizing microbial community. Linking the performance of the RBCs (measured by ammonia removal) to the ammonia oxidizing microbial community would help determine which members of this community have important roles in the ammonia removal process. The activity of the nitrifiers could be further assessed with differential inhibitor and isotope labelling experiments. Understanding the microorganisms that are present within water treatment systems is an important step towards optimizing operational practices for improved effluent quality in municipal, industrial, and aquaculture-associated water treatment systems.

## Acknowledgements

We thank the operators and staff at the Guelph WWTP for access to their facility and for providing plant data and the Joint Genome Institute for sequencing support. We also thank L. H. Bergstrand for assistance in utilizing the BackBLAST pipeline. This research was supported by Discovery Grants to LAH, ACD, WJP, and JDN from the Natural Sciences and Engineering Research Council of Canada (NSERC).

## References

1. Daims H, Lücker S, Wagner M. A new perspective on microbes formerly known as nitrite-oxidizing bacteria. Trends Microbiol 2016; 24: 699–712.

2. Könneke M, Bernhard AE, de la Torre JR, Walker CB, Waterbury JB, Stahl DA. Isolation of an autotrophic ammonia-oxidizing marine archaeon. Nature 2005; 437: 543–546.

3. Purkhold U, Pommerening-Röser A, Juretschko S, Schmid MC, Koops HP, Wagner M. Phylogeny of all recognized species of ammonia oxidizers based on comparative 16S rRNA and *amoA* sequence analysis: Implications for molecular diversity surveys. Appl Environ Microbiol 2000; 66: 5368–5382.

4. Costa E, Pérez J, Kreft J-U. Why is metabolic labour divided in nitrification? Trends Microbiol 2006; 14: 213–219.

5. Daims H, Lebedeva E V., Pjevac P, Han P, Herbold C, Albertsen M, et al. Complete nitrification by *Nitrospira* bacteria. Nature 2015; 528: 504–509.

6. van Kessel MAHJ, Speth DR, Albertsen M, Nielsen PH, Op den Camp HJM, Kartal B, et al. Complete nitrification by a single microorganism. Nature 2015; 528: 555–559.

7. Pjevac P, Schauberger C, Poghosyan L, Herbold CW, Kessel MA van, Daebeler A, et al. *AmoA*-targeted polymerase chain reaction primers for the specific detection and quantification of comammox *Nitrospira* in the environment. Front Microbiol 2017; 8: 1508.

8. Pinto AJ, Marcus DN, Ijaz Z, Bautista-de los Santos QM, Dick GJ, Raskin L. Metagenomic evidence for the presence of comammox Nitrospira-like bacteria in a drinking water system. mSphere 2015; 1: e00054–15.

9. Palomo A, Fowler SJ, Gülay A, Rasmussen S, Sicheritz-Ponten T, Smets BF. Metagenomic analysis of rapid gravity sand filter microbial communities suggests novel physiology of *Nitrospira* spp. ISME J 2016; 10: 2569–2581.

10. Wang Y, Ma L, Mao Y, Jiang X, Xia Y, Yu K, et al. Comammox in drinking water systems. Water Res 2017; 116: 332–341.

11. Palomo A, Dechesne A, Smets BF. Genomic profiling of *Nitrospira* species reveals ecological success of comammox *Nitrospira*. bioRxiv 2019; 612226.

12. Zhao Z, Huang G, Wang M, Zhou N, He S, Dang C, et al. Abundance and community composition of comammox bacteria in different ecosystems by a universal primer set. Sci Total Environ 2019; 691: 146–155.

13. Fowler SJ, Palomo A, Dechesne A, Mines PD, Smets BF. Comammox *Nitrospira* are abundant ammonia oxidizers in diverse groundwater-fed rapid sand filter communities. Environ Microbiol 2018; 20: 1002–1015.

14. Xia F, Wang J-G, Zhu T, Zou B, Rhee S-K, Quan Z-X. Ubiquity and diversity of complete ammonia oxidizers (comammox). Appl Environ Microbiol 2018; 84: e01390–18.

15. Beach NK, Noguera DR. Design and assessment of species-level qPCR primers targeting comammox. Front Microbiol 2019; 10: 36.

16. Annavajhala MK, Kapoor V, Santo-Domingo J, Chandran K. Comammox functionality identified in diverse engineered biological wastewater treatment systems. Environ Sci Technol Lett 2018; 5: 110–116.

17. Roots P, Wang Y, Rosenthal AF, Griffin JS, Sabba F, Petrovich M, et al. Comammox *Nitrospira* are the dominant ammonia oxidizers in a mainstream low dissolved oxygen nitrification reactor. Water Res 2019; 157: 396–405.

18. Cotto I, Dai Z, Huo L, Anderson CL, Vilardi KJ, Ijaz U, et al. Long solids retention times and attached growth phase favor prevalence of comammox bacteria in nitrogen removal systems. Water Res 2020; 169: 115268.

19. Chao Y, Mao Y, Yu K, Zhang T. Novel nitrifiers and comammox in a full-scale hybrid biofilm and activated sludge reactor revealed by metagenomic approach. Appl Microbiol Biotechnol 2016; 100: 8225–8237.

20. Fan X-Y, Gao J-F, Pan K-L, Li D-C, Dai H-H. Temporal dynamics of bacterial communities and predicted nitrogen metabolism genes in a full-scale wastewater treatment plant. RSC Adv 2017; 7: 56317–56327.

21. Pan KL, Gao JF, Fan XY, Li DC, Dai HH. The more important role of archaea than bacteria in nitrification of wastewater treatment plants in cold season despite their numerical relationships. Water Res 2018; 145: 552–561.

22. Wang M, Huang G, Zhao Z, Dang C, Liu W, Zheng M. Newly designed primer pair revealed dominant and diverse comammox *amoA* gene in full-scale wastewater treatment plants. Bioresour Technol 2018; 270: 580–587.

23. Zheng M, Wang M, Zhao Z, Zhou N, He S, Liu S, et al. Transcriptional activity and diversity of comammox bacteria as a previously overlooked ammonia oxidizing prokaryote in full-scale wastewater treatment plants. Sci Total Environ 2019; 656: 717– 722.

24. Yang Y, Pan J, Zhou Z, Wu J, Liu Y, Lin J-G, et al. Complex microbial nitrogen-cycling networks in three distinct anammox-inoculated wastewater treatment systems. Water Res 2020; 168: 115142.

25. Gonzalez-Martinez A, Rodriguez-Sanchez A, van Loosdrecht MCM, Gonzalez-Lopez J, Vahala R. Detection of comammox bacteria in full-scale wastewater treatment bioreactors using tag-454-pyrosequencing. Environ Sci Pollut Res 2016; 23: 25501–25511.

26. Fan X-Y, Gao J-F, Pan K-L, Li D-C, Zhang L-F, Wang S-J. Shifts in bacterial community composition and abundance of nitrifiers during aerobic granulation in two nitrifying sequencing batch reactors. Bioresour Technol 2018; 251: 99–107.

27. Camejo PY, Santo Domingo J, McMahon KD, Noguera DR. Genome-enabled insights into the ecophysiology of the comammox bacterium “*Candidatus* Nitrospira nitrosa”. mSystems 2017; 2: e00059–17.

28. Metch JW, Wang H, Ma Y, Miller JH, Vikesland PJ, Bott C, et al. Insights gained into activated sludge nitrification through structural and functional profiling of microbial community response to starvation stress. Environ Sci Water Res Technol 2019; 5: 884–896.

29. Sauder LA, Peterse F, Schouten S, Neufeld JD. Low-ammonia niche of ammonia-oxidizing archaea in rotating biological contactors of a municipal wastewater treatment plant. Environ Microbiol 2012; 14: 2589–2600.

30. Sauder LA, Albertsen M, Engel K, Schwarz J, Nielsen PH, Wagner M, et al. Cultivation and characterization of *Candidatus* Nitrosocosmicus exaquare, an ammonia-oxidizing archaeon from a municipal wastewater treatment system. ISME J 2017; 11: 1142–1157.

31. Holmes RM, Aminot A, Kérouel R, Hooker BA, Peterson BJ. A simple and precise method for measuring ammonium in marine and freshwater ecosystems. Can J Fish Aquat Sci 1999; 56: 1801–1808.

32. Poulin P, Pelletier É. Determination of ammonium using a microplate-based fluorometric technique. Talanta 2007; 71: 1500–1506.

33. Miranda KM, Espey MG, Wink DA. A rapid, simple spectrophotometric method for simultaneous detection of nitrate and nitrite. Nitric Oxide 2001; 5: 62–71.

34. Ochsenreiter T, Selezi D, Quaiser A, Bonch-Osmolovskaya L, Schleper C. Diversity and abundance of Crenarchaeota in terrestrial habitats studied by 16S RNA surveys and real time PCR. Environ Microbiol 2003; 5: 787–797.

35. Muyzer G, de Waal EC, Uitterlinden AG. Profiling of complex microbial populations by denaturing gradient gel electrophoresis analysis of polymerase chain reaction-amplified genes coding for 16S rRNA. Appl Environ Microbiol 1993; 59: 695–700.

36. Rotthauwe JH, Witzel KP, Liesack W. The ammonia monooxygenase structural gene *amoA* as a functional marker: Molecular fine-scale analysis of natural ammonia-oxidizing populations. Appl Environ Microbiol 1997; 63: 4704–4712.

37. Parada AE, Needham DM, Fuhrman JA. Every base matters: assessing small subunit rRNA primers for marine microbiomes with mock communities, time series and global field samples. Environ Microbiol 2016; 18: 1403–1414.

38. Quince C, Lanzen A, Davenport RJ, Turnbaugh PJ. Removing noise from pyrosequenced amplicons. BMC Bioinformatics 2011; 12: 38.

39. Bolyen E, Rideout JR, Dillon MR, Bokulich NA, Abnet C, Al-Ghalith GA, et al. QIIME 2: Reproducible, interactive, scalable, and extensible microbiome data science. PeerJ Prepr 2018; 6: e27295v2.

40. Martin M. Cutadapt removes adapter sequences from high-throughput sequencing reads. EMBnet.journal 2011; 17: 10.

41. Callahan BJ, McMurdie PJ, Rosen MJ, Han AW, Johnson AJA, Holmes SP. DADA2: High-resolution sample inference from Illumina amplicon data. Nat Methods 2016; 13: 581–583.

42. Quast C, Pruesse E, Yilmaz P, Gerken J, Schweer T, Yarza P, et al. The SILVA ribosomal RNA gene database project: improved data processing and web-based tools. Nucleic Acids Res 2012; 41: D590–D596.

43. Pedregosa F, Varoquaux G, Gramfort A, Michel V, Thirion B, Grisel O, et al. Scikit-learn: Machine learning in Python. J Mach Learn Res 2011; 12: 2825–2830.

44. Schubert M, Lindgreen S, Orlando L. AdapterRemoval v2: rapid adapter trimming, identification, and read merging. BMC Res Notes 2016; 9: 88.

45. Andrews S. FastQC: a quality control tool for high throughput sequence data. http://www.bioinformatics.babraham.ac.uk/projects/fastqc.

46. Ewels P, Magnusson M, Lundin S, Käller M. MultiQC: summarize analysis results for multiple tools and samples in a single report. Bioinformatics 2016; 32: 3047–3048.

47. Kim D, Hahn AS, Wu SJ, Hanson NW, Konwar KM, Hallam SJ. FragGeneScan-plus for scalable high-throughput short-read open reading frame prediction. 2015 IEEE Conf. Comput. Intell. Bioinforma. Comput. Biol. CIBCB 2015. 2015. IEEE, pp 1–8.

48. Fish JA, Chai B, Wang Q, Sun Y, Brown CT, Tiedje JM, et al. FunGene: the functional gene pipeline and repository. Front Microbiol 2013; 4: 291.

49. Petrenko P, Lobb B, Kurtz DA, Neufeld JD, Doxey AC. MetAnnotate: function-specific taxonomic profiling and comparison of metagenomes. BMC Biol 2015; 13: 92.

50. White RAI, Brown J, Colby S, Overall CC, Lee J-Y, Zucker J, et al. ATLAS (Automatic Tool for Local Assembly Structures)-a comprehensive infrastructure for assembly, annotation, and genomic binning of metagenomic and metatranscriptomic data. PeerJ Prepr 2017; 5: e2843v1.

51. Kieser S, Brown J, Zdobnov EM, Trajkovski M, McCue LA. ATLAS: a Snakemake workflow for assembly, annotation, and genomic binning of metagenome sequence data. bioRxiv 2019; 737528.

52. Parks DH, Chuvochina M, Waite DW, Rinke C, Skarshewski A, Chaumeil P-A, et al. A standardized bacterial taxonomy based on genome phylogeny substantially revises the tree of life. Nat Biotechnol 2018; 36: 996–1004.

53. Chaumeil P-A, Hugenholtz P, Parks DH. GTDB-Tk: a toolkit to classify genomes with the Genome Taxonomy Database. Bioinformatics 2019; doi: 10.1093/bioinformatics/btz848.

54. Jain C, Rodriguez-R LM, Phillippy AM, Konstantinidis KT, Aluru S. High throughput ANI analysis of 90K prokaryotic genomes reveals clear species boundaries. Nat Commun 2018; 9: 5114.

55. Camacho C, Coulouris G, Avagyan V, Ma N, Papadopoulos J, Bealer K, et al. BLAST+: architecture and applications. BMC Bioinformatics 2009; 10: 421.

56. Bergstrand LH, Cardenas E, Holert J, van Hamme JD, Mohn WW. Delineation of steroid-degrading microorganisms through comparative genomic analysis. MBio 2016; 7: e00166.

57. Nguyen LT, Schmidt HA, Von Haeseler A, Minh BQ. IQ-TREE: A fast and effective stochastic algorithm for estimating maximum-likelihood phylogenies. Mol Biol Evol 2015; 32: 268–274.

58. Edgar RC. MUSCLE: Multiple sequence alignment with high accuracy and high throughput. Nucleic Acids Res 2004; 32: 1792–1797.

59. Pester M, Maixner F, Berry D, Rattei T, Koch H, Lücker S, et al. *NxrB* encoding the beta subunit of nitrite oxidoreductase as functional and phylogenetic marker for nitrite-oxidizing *Nitrospira*. Environ Microbiol 2014; 16: 3055–3071.

60. Dahllof I, Baillie H, Kjelleberg S. *rpoB*-based microbial community analysis avoids limitations inherent in 16S rRNA gene intraspecies heterogeneity. Appl Environ Microbiol 2000; 66: 3376–3380.

61. Bartleme RP, Mclellan SL, Newton RJ. Freshwater recirculating aquaculture system operations drive biofilter bacterial community shifts around a stable nitrifying consortium of ammonia-oxidizing archaea and comammox *Nitrospira*. Front Microbiol 2017; 8: 101.

62. Park H-D, Wells GF, Bae H, Criddle CS, Francis CA. Occurrence of ammonia-oxidizing archaea in wastewater treatment plant bioreactors. Appl Environ Microbiol 2006; 72: 5643–5647.

63. Kits KD, Sedlacek CJ, Lebedeva E V., Han P, Bulaev A, Pjevac P, et al. Kinetic analysis of a complete nitrifier reveals an oligotrophic lifestyle. Nature 2017; 549: 269–272.

64. Bai Y, Sun Q, Wen D, Tang X. Abundance of ammonia-oxidizing bacteria and archaea in industrial and domestic wastewater treatment systems. FEMS Microbiol Ecol 2012; 80: 323–330.

65. Roy D, McEvoy J, Blonigen M, Amundson M, Khan E. Seasonal variation and ex-situ nitrification activity of ammonia oxidizing archaea in biofilm based wastewater treatment processes. Bioresour Technol 2017; 244: 850–859.

66. Gao J-F, Luo X, Wu G-X, Li T, Peng Y-Z. Quantitative analyses of the composition and abundance of ammonia-oxidizing archaea and ammonia-oxidizing bacteria in eight full-scale biological wastewater treatment plants. Bioresour Technol 2013; 138: 285–296.

67. Zhang T, Ye L, Tong AHY, Shao MF, Lok S. Ammonia-oxidizing archaea and ammonia-oxidizing bacteria in six full-scale wastewater treatment bioreactors. Appl Microbiol Biotechnol 2011; 91: 1215–1225.

68. Limpiyakorn T, Sonthiphand P, Rongsayamanont C, Polprasert C. Abundance of *amoA* genes of ammonia-oxidizing archaea and bacteria in activated sludge of full-scale wastewater treatment plants. Bioresour Technol 2011; 102: 3694–3701.

69. Lawson CE, Lücker S. Complete ammonia oxidation: an important control on nitrification in engineered ecosystems? Curr Opin Biotechnol 2018; 50: 158–165.

70. Palomo A, Pedersen AG, Fowler SJ, Dechesne A, Sicheritz-Pontén T, Smets BF. Comparative genomics sheds light on niche differentiation and the evolutionary history of comammox *Nitrospira*. ISME J 2018; 12: 1779–1793.

71. Koch H, Lücker S, Albertsen M, Kitzinger K, Herbold C, Spieck E, et al. Expanded metabolic versatility of ubiquitous nitrite-oxidizing bacteria from the genus *Nitrospira*. Proc Natl Acad Sci U S A 2015; 112: 11371–11376.

72. Lücker S, Wagner M, Maixner F, Pelletier E, Koch H, Vacherie B, et al. A *Nitrospira* metagenome illuminates the physiology and evolution of globally important nitrite-oxidizing bacteria. Proc Natl Acad Sci U S A 2010; 107: 13479–13484.

73. Ushiki N, Fujitani H, Shimada Y, Morohoshi T, Sekiguchi Y, Tsuneda S. Genomic analysis of two phylogenetically distinct *Nitrospira* species reveals their genomic plasticity and functional diversity. Front Microbiol 2018; 8: 2637.

74. Poghosyan L, Koch H, Lavy A, Frank J, Kessel MAHJ, Jetten MSM, et al. Metagenomic recovery of two distinct comammox *Nitrospira* from the terrestrial subsurface. Environ Microbiol 2019; E-pub DOI 10.1111/1462-2920.14691.

75. Coppi M V. The hydrogenases of *Geobacter sulfurreducens*: a comparative genomic perspective. Microbiology 2005; 151: 1239–1254.

76. Koch H, Galushko A, Albertsen M, Schintlmeister A, Gruber-Dorninger C, Lücker S, et al. Growth of nitrite-oxidizing bacteria by aerobic hydrogen oxidation. Science 2014; 345: 1052–1054.

77. Palatinszky M, Herbold C, Jehmlich N, Pogoda M, Han P, von Bergen M, et al. Cyanate as an energy source for nitrifiers. Nature 2015; 524: 105–108.

78. Koch H, van Kessel MAHJ, Lücker S. Complete nitrification: insights into the ecophysiology of comammox *Nitrospira*. Appl Microbiol Biotechnol 2019; 103: 177–189.

79. Walsh MA, Otwinowski Z, Perrakis A, Anderson PM, Joachimiak A. Structure of cyanase reveals that a novel dimeric and decameric arrangement of subunits is required for formation of the enzyme active site. Structure 2000; 8: 505–14.

80. Marier JR, Rose D. Determination of cyanate, and a study of its accumulation in aqueous solutions of urea. Anal Biochem 1964; 7: 304–314.

81. Dirnhuber P, Schütz F. The isomeric transformation of urea into ammonium cyanate in aqueous solutions. Biochem J 1948; 42: 628–32.

82. Anderson PM, Sung Y, Fuchs JA. The cyanase operon and cyanate metabolism. FEMS Microbiol Lett 1990; 87: 247–252.

83. Purcarea C, Ahuja A, Lu T, Kovari L, Guy HI, Evans DR. *Aquifex aeolicus* aspartate transcarbamoylase, an enzyme specialized for the efficient utilization of unstable carbamoyl phosphate at elevated temperature. J Biol Chem 2003; 278: 52924–34.

84. Kamennaya NA, Post AF. Characterization of cyanate metabolism in marine *Synechococcus* and *Prochlorococcus* spp. Appl Environ Microbiol 2011; 77: 291–301.

85. Muñoz-Centeno MC, Paneque A, Cejudo FJ. Cyanate is transported by the nitrate permease in *Azotobacter chroococcum*. FEMS Microbiol Lett 1996; 137: 91–94.

86. Lehtovirta-Morley LE. Ammonia oxidation: Ecology, physiology, biochemistry and why they must all come together. FEMS Microbiol Lett 2018; 365: fny058.

87. Prosser JI, Nicol GW. Archaeal and bacterial ammonia-oxidisers in soil: The quest for niche specialisation and differentiation. Trends Microbiol 2012; 20: 523–531.

88. Probst AJ, Auerbach AK, Moissl-Eichinger C. Archaea on human skin. PLoS One 2013; 8: e65388.

89. Daims H, Nielsen JL, Nielsen PH, Schleifer KH, Wagner M. In situ characterization of *Nitrospira*-like nitrite-oxidizing bacteria active in wastewater treatment plants. Appl Environ Microbiol 2001; 67: 5273–5284.

90. Gruber-Dorninger C, Pester M, Kitzinger K, Savio DF, Loy A, Rattei T, et al. Functionally relevant diversity of closely related *Nitrospira* in activated sludge. ISME J 2015; 9: 643–655.

