## Supplementary Information for "High functional diversity among *Nitrospira* populations that dominate rotating biological contactor microbial communities in a municipal wastewater treatment plant"

Running title: Comammox bacteria dominate wastewater biofilm

Keywords: complete ammonia oxidation, comammox, ammonia-oxidizing archaea, wastewater, nitrification, biofilm, cyanate, metagenomics

**Supplemental materials and methods**

### Water chemistry

Ammonium was measured fluorometrically using orthophthaldialdehyde (OPA) reagent [1] according to the method by Poulin and Pelletier [2], with minor modifications. Volumes of 100 µL of sample and 200 µL of OPA working reagent were added to a 96-well opaque flat-bottomed plate. Plates were incubated for four hours in the dark before being measured. Nitrite and nitrate concentrations were assessed colorimetrically using the Greiss reagent as described elsewhere [3]. All samples were measured as technical duplicates at 360 nm excitation, 465 nm emission (ammonium), and 550 nm (nitrite/nitrate) using a FilterMax F5 Multi-Mode Microplate reader (Molecular Devices, San Jose, CA, USA). Measurements represent total ammonium as nitrogen, and nitrite and nitrate as nitrogen. The pH of water samples was measured with a DELTA 320 pH meter (Mettler Toledo, Mississauga, ON, Canada).

### Quantitative PCR

Each 10 µL PCR contained 1X iQ SYBR Green Supermix (Bio-Rad; containing 50 U/ml iTaq DNA polymerase, 0.4 mM dNTPs, 6 mM MgCl_2_, 100 mM KCl, 40 mM Tris-HCl, 20 nM fluorescein, stabilizers, and SYBR Green I dye), primers (see below for concentrations), 0.5 µg bovine serum albumin, and 2-20 ng of DNA as template. Primers were used at a final concentration of 0.2 µM, except for the comaA primers, which were used at 0.5 µM. Conditions for qPCR of bacterial 16S rRNA, AOA 16S rRNA, and AOB *amoA* genes were 95°C for 3 minutes, followed by 35 cycles of 95°C for 30 s, 55°C for 30 s, and 72°C for 30 s, with a fluorescence reading at each step. Melt curve analysis was performed from 65-95°C in increments of 0.5°C. The qPCR conditions for comammox *Nitrospira* *amoA* genes were the same except that the annealing temperature was 52°C and the extension time at 72°C was 1 minute. Standard curves were generated using 10-fold dilutions of standards that were made using the same primer pairs used for qPCR. For comammox *Nitrospira* *amoA* genes, template DNA for the standard curve was extracted from the rotating biological contactor (RBC) biofilm*.* For AOB *amoA* genes and AOA 16S rRNA genes, template DNA for the standard curve was extracted from cultures of *Nitrosomonas europaea* and *Candidatus* Nitrosocosmicus exaquare, respectively. Extracted DNA from *Escherichia coli* was used to prepare standards for bacterial 16S rRNA gene qPCR. Gene copies were calculated based on the amount of DNA present in the original extractions but were not normalized further to account for copy number.

### Assembly and binning of metagenomic reads

Within ATLAS, quality control was performed using the BBTools suite of utilities, version 37.78 (BBMap – Bushnell B. – [sourceforge.net/projects/bbmap/](http://sourceforge.net/projects/bbmap/)). Individual assemblies and several binning strategies were performed (Table S3). First, samples were assembled using SPADES 3.12.0 [4]. Two different binning tools, MaxBin2 version 2.2.4 [5] and Metabat2 version 2.12.1 [6], were used, and DASTool version 1.1.1 [7] was used to select the best bins. Bin completeness and contamination were assessed using CheckM version 1.0.7 [8]. Genome bins from assemblies were subsequently dereplicated to find identical genomes using dRep version 2.2.2 [9], using a 99% threshold for clustering. Only bins that met the threshold of 75% completeness and 15% contamination were included in the dereplicated set. Lastly, predicted proteins from contigs were clustered using the linclust module of MMseqs2, version 3.be8f6 [10]. All settings used in the ATLAS pipeline are available in the run configuration file, Supplementary File 3. Note that the "--only-assembler" flag in SPADES had to be enabled for two samples, SE8 and FNE8, to work around a known software error during the pre-assembly error correction step of SPADES.

### Analysis of genome bins

The *Nitrospira* bins were further searched for genes involved in nitrogen cycling using reciprocal BLASTP [11] of known genes in four reference *Nitrospira* genomes (*Ca.* N. inopinata, *Nitrospira moscoviensis*, *Nitrospira lenta,* and *Nitrospira japonica*). The searches were performed using the BackBLAST pipeline [12], development version 2.0.0-alpha2 (doi:[10.5281/zenodo.3465955](https://doi.org/10.5281/zenodo.3465955)) and the e-value threshold and identity cutoff were manually optimized to 1e-30 and 40%, respectively, based on expected hits to reference genomes. Additionally, as a new default setting compared to the previous release of BackBLAST (v1.0), a soft mask was applied during the database search phase of BLASTp to improve ortholog detection [13]. To prevent the incorrect exclusion of true orthologs among AmoC proteins, the AmoC3 protein sequence (WP_062483313.1) was removed from the reference proteome of *Ca.* N. inopinata during the BLASTP search due to its close sequence identity to the query AmoC1 protein (WP_062484140.1), which interfered with reciprocal BLASTP.

A concatenated core protein phylogeny was created to assess the phylogenetic placement of the *Nitrospira* genome bins. Reference genomes were downloaded from the NCBI database and a core set of 74 core bacterial proteins ('Bacteria.hmm' in GToTree) were identified, aligned, concatenated, and used for phylogenetic tree construction from these and the *Nitrospira* genome bins via the GToTree pipeline, version 1.1.10 [14], with default settings. The maximum likelihood phylogeny was computed using IQ-TREE version 1.6.9 [15] with 1000 bootstrap replicates and the LG+F+R10 evolutionary model as selected by IQ-TREE's ModelFinder module [16] .

### Functional gene phylogenies

The clustered sequence set of predicted protein sequences was searched with the "amoA_AOB" profile hidden Markov model (HMM) from the FunGene database [17] using hmmsearch version 3.1b2 [18] with an e-value cutoff of 1e-10 to identify putative AmoA sequences. Identified hits were queried against the NCBI RefSeq database via BLASTP [11] and those with closest database matches to the *Nitrospirota* phylum were used for downstream analysis. Using MUSCLE [19] within MEGA7 [20], sequences of identified proteins were aligned, along with other AmoA sequences from enrichment cultures and environmental surveys retrieved from the NCBI database. Phylogenetic analysis was inferred using maximum likelihood analysis with 500 bootstrap replicates, based on the Le Gascuel evolutionary model [21], using MEGA7.

Cyanase protein sequences (CynS) identified via BackBLAST were further analyzed phylogenetically. Reference annotated CynS sequences were identified from the SwissProt database and downloaded from RefSeq for consistency with other analytical steps. These references were aligned to the CynS sequences identified in *Nitrospira* genomes using Clustal Omega version 1.2.3 [22, 23], and the alignment was used to build a maximum likelihood phylogeny using IQ-TREE version 1.6.11 with 1000 rapid bootstraps [15]. The LG+I+G4 evolutionary model was selected based on the ModelFinder module of IQ-TREE [16]. The two annotated archaeal reference sequences of CynS were chosen as the outgroup for the phylogeny as shown previously [24]. Subsequently, all genome bins in the metagenomic data were searched for the presence of CynS using hmmsearch version 3.1b2 [18]. The Pfam profile HMM PF02560, which covers the unique C-terminal domain of CynS, was used to perform the search with an e-value cutoff of 1e-20. Read mapping statistics for genome bins containing CynS, along with the CynS multiple sequence alignment and phylogeny, were plotted using a custom R script.

**References**

1.	Holmes RM, Aminot A, Kérouel R, Hooker BA, Peterson BJ. A simple and precise method for measuring ammonium in marine and freshwater ecosystems. *Can J Fish Aquat Sci* 1999; **56**: 1801–1808.

2.	Poulin P, Pelletier É. Determination of ammonium using a microplate-based fluorometric technique. *Talanta* 2007; **71**: 1500–1506.

3.	Miranda KM, Espey MG, Wink DA. A rapid, simple spectrophotometric method for simultaneous detection of nitrate and nitrite. *Nitric Oxide* 2001; **5**: 62–71.

4.	Nurk S, Meleshko D, Korobeynikov A, Pevzner PA. metaSPAdes: a new versatile metagenomic assembler. *Genome Res* 2017; **27**: 824–834.

5.	Wu Y-W, Simmons BA, Singer SW. MaxBin 2.0: an automated binning algorithm to recover genomes from multiple metagenomic datasets. *Bioinformatics* 2016; **32**: 605–607.

6.	Kang DD, Froula J, Egan R, Wang Z. MetaBAT, an efficient tool for accurately reconstructing single genomes from complex microbial communities. *PeerJ* 2015; **3**: e1165.

7.	Sieber CMK, Probst AJ, Sharrar A, Thomas BC, Hess M, Tringe SG, et al. Recovery of genomes from metagenomes via a dereplication, aggregation and scoring strategy. *Nat Microbiol* 2018; **3**: 836–843.

8.	Parks DH, Imelfort M, Skennerton CT, Hugenholtz P, Tyson GW. CheckM: assessing the quality of microbial genomes recovered from isolates, single cells, and metagenomes. *Genome Res* 2015; **25**: 1043–1055.

9.	Olm MR, Brown CT, Brooks B, Banfield JF. dRep: a tool for fast and accurate genomic comparisons that enables improved genome recovery from metagenomes through de-replication. *ISME J* 2017; **11**: 2864–2868.

10.	Steinegger M, Söding J. Clustering huge protein sequence sets in linear time. *Nat Commun* 2018; **9**: 2542.

11.	Camacho C, Coulouris G, Avagyan V, Ma N, Papadopoulos J, Bealer K, et al. BLAST+: architecture and applications. *BMC Bioinformatics* 2009; **10**: 421.

12.	Bergstrand LH, Cardenas E, Holert J, van Hamme JD, Mohn WW. Delineation of steroid-degrading microorganisms through comparative genomic analysis. *MBio* 2016; **7**: e00166.

13.	Moreno-Hagelsieb G and Latimer K. Choosing BLAST options for better detection of orthologs as reciprocal best hits. *Bioinformatics* 2008; 24: 319–324.

14.	Lee MD. GToTree: a user-friendly workflow for phylogenomics. *Bioinformatics* 2019; 1–3.

15.	Nguyen LT, Schmidt HA, Von Haeseler A, Minh BQ. IQ-TREE: A fast and effective stochastic algorithm for estimating maximum-likelihood phylogenies. *Mol Biol Evol* 2015; **32**: 268–274.

16.	Kalyaanamoorthy S, Minh BQ, Wong TKF, von Haeseler A, Jermiin LS. ModelFinder: fast model selection for accurate phylogenetic estimates. *Nat Methods* 2017; **14**: 587–589.

17.	Fish JA, Chai B, Wang Q, Sun Y, Brown CT, Tiedje JM, et al. FunGene: the functional gene pipeline and repository. *Front Microbiol* 2013; **4**: 291.

18.	Eddy SR. Accelerated Profile HMM Searches. *PLoS Comput Biol* 2011; **7**: e1002195.

19.	Edgar RC. MUSCLE: Multiple sequence alignment with high accuracy and high throughput. *Nucleic Acids Res* 2004; **32**: 1792–1797.

20.	Kumar S, Stecher G, Tamura K. MEGA7: Molecular Evolutionary Genetics Analysis version 7.0 for bigger datasets. *Mol Biol Evol* 2016; **33**: 1870–1874.

21.	Le SQ, Gascuel O. An improved general amino acid replacement matrix. *Mol Biol Evol* 2008; **25**: 1307–1320.

22.	Sievers F, Wilm A, Dineen D, Gibson TJ, Karplus K, Li W, et al. Fast, scalable generation of high‐quality protein multiple sequence alignments using Clustal Omega. *Mol Syst Biol* 2011; **7**: 539.

23.	Sievers F, Higgins DG. Clustal Omega for making accurate alignments of many protein sequences. *Protein Sci* 2018; **27**: 135–145.

24.	Kamennaya NA, Post AF. Characterization of cyanate metabolism in marine *Synechococcus* and *Prochlorococcus* spp. *Appl Environ Microbiol* 2011; **77**: 291–301.

**Supplementary Tables**

**Table S1** Rotating biological contactor (RBC) samples used in this study.

| Abbreviation | RBC stage | RBC train | Month | Year | Study | Analysis |
| --- | --- | --- | --- | --- | --- | --- |
| FNE1 | RBC 1 | NE train | February | 2010 | Sauder et al., 2012 | qPCR, metagenomics, amplicon sequencing |
| FNE8 | RBC 8 | NE train | February | 2010 | Sauder et al., 2012 | qPCR, metagenomics, amplicon sequencing |
| JNE1 | RBC 1 | NE train | June | 2010 | Sauder et al., 2012 | qPCR, metagenomics, amplicon sequencing |
| JNE8 | RBC 8 | NE train | June | 2010 | Sauder et al., 2012 | qPCR, metagenomics, amplicon sequencing |
| SNE1 | RBC 1 | NE train | September | 2010 | Sauder et al., 2012 | qPCR, metagenomics, amplicon sequencing |
| SNE8 | RBC 8 | NE train | September | 2010 | Sauder et al., 2012 | qPCR, metagenomics, amplicon sequencing |
| NE1 | RBC 1 | NE train | October | 2016 | This study | qPCR, metagenomics, amplicon sequencing |
| NE2 | RBC 2 | NE train | October | 2016 | This study | qPCR |
| NE3 | RBC 3 | NE train | October | 2016 | This study | qPCR |
| NE4 | RBC 4 | NE train | October | 2016 | This study | qPCR |
| NE5 | RBC 5 | NE train | October | 2016 | This study | qPCR |
| NE6 | RBC 6 | NE train | October | 2016 | This study | qPCR |
| NE7 | RBC 7 | NE train | October | 2016 | This study | qPCR |
| NE8 | RBC 8 | NE train | October | 2016 | This study | qPCR, metagenomics, amplicon sequencing |
| NW1 | RBC 1 | NW train | October | 2016 | This study | qPCR, metagenomics, amplicon sequencing |
| NW2 | RBC 2 | NW train | October | 2016 | This study | qPCR |
| NW3 | RBC 3 | NW train | October | 2016 | This study | qPCR |
| NW4 | RBC 4 | NW train | October | 2016 | This study | qPCR |
| NW5 | RBC 5 | NW train | October | 2016 | This study | qPCR |
| NW6 | RBC 6 | NW train | October | 2016 | This study | qPCR |
| NW7 | RBC 7 | NW train | October | 2016 | This study | qPCR |
| NW8 | RBC 8 | NW train | October | 2016 | This study | qPCR, metagenomics, amplicon sequencing |
| SE2 | RBC 2 | SE train | October | 2016 | This study | qPCR, metagenomics, amplicon sequencing |
| SE3 | RBC 3 | SE train | October | 2016 | This study | qPCR |
| SE4 | RBC 4 | SE train | October | 2016 | This study | qPCR |
| SE5 | RBC 5 | SE train | October | 2016 | This study | qPCR |
| SE6 | RBC 6 | SE train | October | 2016 | This study | qPCR |
| SE7 | RBC 7 | SE train | October | 2016 | This study | qPCR |
| SE8 | RBC 8 | SE train | October | 2016 | This study | qPCR, metagenomics, amplicon sequencing |
| SW1 | RBC 1 | SW train | October | 2016 | This study | qPCR, metagenomics, amplicon sequencing |
| SW3 | RBC 3 | SW train | October | 2016 | This study | qPCR |
| SW4 | RBC 4 | SW train | October | 2016 | This study | qPCR |
| SW5 | RBC 5 | SW train | October | 2016 | This study | qPCR |
| SW6 | RBC 6 | SW train | October | 2016 | This study | qPCR |
| SW7 | RBC 7 | SW train | October | 2016 | This study | qPCR |
| SW8 | RBC 8 | SW train | October | 2016 | This study | qPCR, metagenomics, amplicon sequencing |

**Table S2** Primers used for qPCR amplifications.

| Target | Primer | Sequence (5' to 3') | Reference |
| --- | --- | --- | --- |
| AOA 16S rRNA gene | 771F | ACGGTGAGGGATGAAAGCT | Ochsenreiter *et al.*, 2003 |
|  | 957R | CGGCGTTGACTCCA ATTG | Ochsenreiter *et al.*, 2003 |
| Bacterial 16S rRNA gene | 341F | CCTACGGGAGGCAGCAG | Muyzer *et al.*, 1993 |
|  | 518R | ATTACCGCGGCTGCTGG | Muyzer *et al.*, 1993 |
| AOB *amoA* | amoA1F | GGGGTTTCTACTGGTGGT | Rotthauwe *et al.*, 1997 |
|  | amoA2R | CCCCTCKGSAAAGCCTTCTTC | Rotthauwe *et al.*, 1997 |
| Comammox *Nitrospira* clade A *amoA* | comaA-244f * | TACAACTGGGTGAACTA | Pjevac *et al.*, 2017 |
|  |  | TATAACTGGGTGAACTA |  |
|  |  | TACAATTGGGTGAACTA |  |
|  |  | TACAACTGGGTCAACTA |  |
|  |  | TACAACTGGGTCAATTA |  |
|  |  | TATAACTGGGTCAATTA |  |
|  | comaA-659r * | AGATCATGGTGCTATG | Pjevac *et al.*, 2017 |
|  |  | AAATCATGGTGCTATG |  |
|  |  | AGATCATGGTGCTGTG |  |
|  |  | AAATCATGGTGCTGTG |  |
|  |  | AGATCATCGTGCTGTG |  |
|  |  | AAATCATCGTGCTGTG |  |
| Comammox *Nitrospira* clade B *amoA* | comaB-244f * | TAYTTCTGGACGTTCTA | Pjevac *et al.*, 2017 |
|  |  | TAYTTCTGGACATTCTA |  |
|  |  | TACTTCTGGACTTTCTA |  |
|  |  | TAYTTCTGGACGTTTTA |  |
|  |  | TAYTTCTGGACATTTTA |  |
|  |  | TACTTCTGGACCTTCTA |  |
|  | comaB-659r * | ARATCCAGACGGTGTG | Pjevac *et al.*, 2017 |
|  |  | ARATCCAAACGGTGTG |  |
|  |  | ARATCCAGACAGTGTG |  |
|  |  | ARATCCAAACAGTGTG |  |
|  |  | AGATCCAGACTGTGTG |  |
|  |  | AGATCCAAACAGTGTG |  |
| *equimolar primer mix of all six primers | |  |  |

**Table S3** Metagenome sequencing and assembly data for rotating biological contactor (RBC) samples.

| Sample | Sequenced read pairs (million) | Read pairs after quality filtering (million) | Assembled reads | Assembled reads (%) | Number of predicted genes | Total size of contigs (Mb) | Number of contigs | Number of scaffolds | L50 | L90 | N50 | N90 | Average G + C content (%) |
| --- | --- | --- | --- | --- | --- | --- | --- | --- | --- | --- | --- | --- | --- |
| FNE1 | 19.6 | 19.6 | 13.9 | 35.4 | 297430 | 257.3 | 80985 | 77575 | 5517 | 1212 | 8409 | 53766 | 60.2 |
| FNE8 | 16.2 | 16.1 | 8.3 | 25.9 | 254136 | 223.4 | 65269 | 62389 | 6991 | 1222 | 5308 | 42485 | 60.9 |
| JNE1 | 20.3 | 20.3 | 12.7 | 31.4 | 398212 | 341.0 | 101976 | 96908 | 6304 | 1233 | 9227 | 66060 | 54.4 |
| JNE8 | 21.6 | 21.6 | 13.9 | 32.4 | 363091 | 311.5 | 95133 | 90192 | 6090 | 1215 | 8574 | 62034 | 52.8 |
| SNE1 | 19.3 | 19.3 | 11.9 | 30.9 | 311133 | 269.3 | 84717 | 81836 | 5765 | 1190 | 7220 | 56514 | 61.4 |
| SNE8 | 21.6 | 21.6 | 11.2 | 26.0 | 401942 | 345.7 | 121415 | 116947 | 4101 | 1155 | 13389 | 85192 | 62.8 |
| NE1 | 17.0 | 17.0 | 10.4 | 30.6 | 291270 | 257.0 | 74472 | 70350 | 6618 | 1252 | 6728 | 48089 | 63.0 |
| NE8 | 11.4 | 11.4 | 4.0 | 17.6 | 169150 | 142.8 | 51200 | 48840 | 3766 | 1183 | 7489 | 36802 | 63.4 |
| NW1 | 20.3 | 20.3 | 12.3 | 30.3 | 380050 | 333.1 | 102953 | 96974 | 5582 | 1224 | 10421 | 68044 | 63.2 |
| NW8 | 10.5 | 10.5 | 3.7 | 17.8 | 152896 | 129.4 | 45676 | 43920 | 3995 | 1165 | 5661 | 32521 | 63.3 |
| SE2 | 20.7 | 20.6 | 12.7 | 30.8 | 341712 | 297.7 | 91461 | 86732 | 5432 | 1245 | 9421 | 60088 | 62.3 |
| SE8 | 16.2 | 16.2 | 7.4 | 23.1 | 253689 | 222.3 | 67483 | 63503 | 5912 | 1225 | 6296 | 44982 | 63.4 |
| SW1 | 19.6 | 19.6 | 10.3 | 26.2 | 361461 | 310.9 | 103644 | 98863 | 4811 | 1178 | 11189 | 70590 | 61.7 |
| SW8 | 18.4 | 18.4 | 9.6 | 26.2 | 317476 | 278.2 | 83699 | 79008 | 5918 | 1240 | 7798 | 55048 | 62.5 |

**Table S4** Water chemistry for October 2016 rotating biological contactor (RBC) samples.

|  | Ammonium | | Nitrite | | Nitrate | |
| --- | --- | --- | --- | --- | --- | --- |
| Sample | Concentration (µM) | Loading (kg/day) * | Concentration (µM) | Loading (kg/day) * | Concentration (µM) | Loading (kg/day) * |
| NE1 | 16.3 | 12.1 | 66.1 | 49.0 | 1778 | 13198 |
| NE8 | 3.6 | 2.7 | 3.3 | 2.4 | 1471 | 10923 |
| NW1 | 14.3 | 10.6 | 31.5 | 23.4 | 682 | 5062 |
| NW8 | 0.2 | 0.2 | 0.9 | 0.7 | 359 | 2662 |
| SE1 | 15.4 | 11.4 | 27.7 | 20.6 | 567 | 4206 |
| SE8 | 1.1 | 0.8 | 6.2 | 4.6 | 633 | 4696 |
| SW1 | 9.3 | 6.9 | 18.4 | 13.7 | 478 | 3548 |
| SW8 | 2.1 | 1.5 | 9.8 | 7.3 | 970 | 7202 |
| * Loading was calculated based on the average flow rate of the Guelph WWTP in 2016 (53000 m³/day). | | | | | | |
| The measured concentration (in mg/L) was converted to kg/L and then was multiplied by the average flow rate (in L/day) | | | | | | |

**Table S5** Quantitative PCR results for rotating biological contactor (RBC) biofilm samples.

|  | Bacterial 16S rRNA | |  |  | Comammox *Nitrospira* *amoA* | |  |  | AOA 16S rRNA | |  | AOB *amoA* | |  | Total AOP |  |  |  |  |  |  |
| --- | --- | --- | --- | --- | --- | --- | --- | --- | --- | --- | --- | --- | --- | --- | --- | --- | --- | --- | --- | --- | --- |
| Biofilm sample | Copies/ng genomic DNA ^†^ | SD |  |  | Copies/ng genomic DNA ^†^ | SD |  |  | Copies/ng genomic DNA ^†^ | SD |  | Copies/ng genomic DNA ^†^ | SD |  | Copies/ng genomic DNA ^†^ | AOA of total AOP (%) |  | AOB of total AOP (%) | AOA of TC (%) |  | AOB of TC (%) |
|  |  |  |  |  |  |  |  |  |  |  |  |  |  |  |  |  | Comammox *Nitrospira* of total AOP (%) |  |  | Comammox *Nitrospira* of TC (%) |  |
| FNE1 | 89448 | 13334 |  |  | 12754 | 1892 |  |  | 392* | 65 |  | 2315* | 25 |  | 15461 | 2.5 | 82.5 | 15 | 0.4 | 14.3 | 2.6 |
| FNE8 | 81515 | 6044 |  |  | 6517 | 414 |  |  | 2257* | 144 |  | 1216* | 35 |  | 9990 | 22.6 | 65.2 | 12.2 | 2.7 | 8 | 1.5 |
| JNE1 | 77613 | 5965 |  |  | 5180 | 2102 |  |  | 148 | 19 |  | 1755 | 206 |  | 7083 | 2.1 | 73.1 | 24.8 | 0.2 | 6.7 | 2.3 |
| JNE8 | 95455 | 1332 |  |  | 4752 | 1588 |  |  | 1063 | 249 |  | 1131 | 86 |  | 6947 | 15.3 | 68.4 | 16.3 | 1.1 | 5 | 1.2 |
| SNE1 | 133595 | 19159 |  |  | 20150* | 513 |  |  | 2150 | 498 |  | 2387 | 583 |  | 24687 | 8.7 | 81.6 | 9.7 | 1.6 | 15.1 | 1.8 |
| SNE8 | 117389 | 41914 |  |  | 2554* | 706 |  |  | 3863 | 139 |  | 508 | 118 |  | 6925 | 55.8 | 36.9 | 7.3 | 3.2 | 2.2 | 0.4 |
| NE1 | 99901 | 9883 |  |  | 5824 | 1094 |  |  | 4500 | 710 |  | 209 | 25 |  | 10532 | 42.7 | 55.3 | 2 | 4.3 | 5.8 | 0.2 |
| NE8 | 75883 | 0^††^ |  |  | 1348 | 80 |  |  | 1783 | 174 |  | 27 | 5 |  | 3158 | 56.5 | 42.7 | 0.9 | 2.3 | 1.8 | 0 |
| NW1 | 103216 | 18345 |  |  | 6334 | 1302 |  |  | 4437* | 144 |  | 297* | 24 |  | 11068 | 40.1 | 57.2 | 2.7 | 4.1 | 6.1 | 0.3 |
| NW8 | 93546 | 15074 |  |  | 1710 | 394 |  |  | 2099* | 376 |  | 21* | 0 |  | 3829 | 54.8 | 44.6 | 0.5 | 2.2 | 1.8 | 0 |
| SE2 | 119854 | 3149 |  |  | 7978 | 21 |  |  | 6868 | 280 |  | 161 | 19 |  | 15007 | 45.8 | 53.2 | 1.1 | 5.4 | 6.7 | 0.1 |
| SE8 | 69612 | 12123 |  |  | 3804 | 1315 |  |  | 3051 | 155 |  | 99 | 22 |  | 6954 | 43.9 | 54.7 | 1.4 | 4.2 | 5.5 | 0.1 |
| SW1 | 109594 | 12893 |  |  | 21264 | 3460 |  |  | 4629* | 215 |  | 97 | 12 |  | 25989 | 17.8 | 81.8 | 0.4 | 4.1 | 19.4 | 0.1 |
| SW8 | 95913 | 14632 |  |  | 9779 | 61 |  |  | 3483* | 141 |  | 60 | 4 |  | 13322 | 26.1 | 73.4 | 0.5 | 3.5 | 10.2 | 0.1 |

† gene copies not standardized by gene copy number

†† no technical duplicate available

* significant difference between RBC 1 and RBC 8 in the same train at the same time point (Welch’s t-test, *p*<0.05)

SD, standard deviation of technical duplicates

AOP, ammonia-oxidizing prokaryotes (AOA plus AOB plus comammox *Nitrospira*)

TC, total community (bacterial plus thaumarchaeotal 16S rRNA genes)

**Table S6** Bin statistics and classification of *Nitrospira* metagenome assembled genomes (MAGs).

| MAG ID | Completeness (%) | Contamination (%) | Genome size (Mbp) | Number of contigs | Contig N50 | GC content (%) | *Nitrospira* lineage |
| --- | --- | --- | --- | --- | --- | --- | --- |
| RBC001 | 94.9 | 2.7 | 3.8 | 227 | 32534 | 54.9 | Clade A comammox |
| RBC003 | 94.0 | 1.8 | 4.3 | 295 | 24838 | 50.1 | Sublineage IV |
| RBC021 | 78.5 | 1.3 | 3.6 | 460 | 11365 | 54.6 | Clade A comammox |
| RBC026 | 75.1 | 3.2 | 2.9 | 567 | 7228 | 58.9 | Sublineage I |
| RBC035 | 94.9 | 2.3 | 4.5 | 473 | 14062 | 55.0 | Clade A comammox |
| RBC042 | 89.6 | 4.8 | 3.4 | 469 | 11565 | 54.7 | Clade A comammox |
| RBC044 | 89.9 | 0.9 | 3.6 | 343 | 15917 | 54.7 | Clade A comammox |
| RBC047 | 75.7 | 3.5 | 3.0 | 542 | 7102 | 54.5 | Clade A comammox |
| RBC048 | 94.9 | 2.7 | 3.9 | 174 | 38558 | 58.6 | Sublineage I |
| RBC069 | 95.9 | 2.7 | 4.5 | 87 | 78382 | 54.5 | Clade A comammox |
| RBC073 | 91.3 | 0.0 | 4.0 | 64 | 161412 | 59.5 | Sublineage I |
| RBC083 | 82.4 | 1.9 | 2.7 | 212 | 21077 | 55.0 | Clade A comammox |
| RBC085 | 92.1 | 1.8 | 3.7 | 250 | 25628 | 59.0 | Sublineage I |
| RBC093 | 94.9 | 2.8 | 3.8 | 58 | 116502 | 55.0 | Clade A comammox |
| RBC100 | 94.0 | 2.3 | 4.3 | 233 | 32473 | 55.2 | Clade A comammox |

**Supplementary Figures**

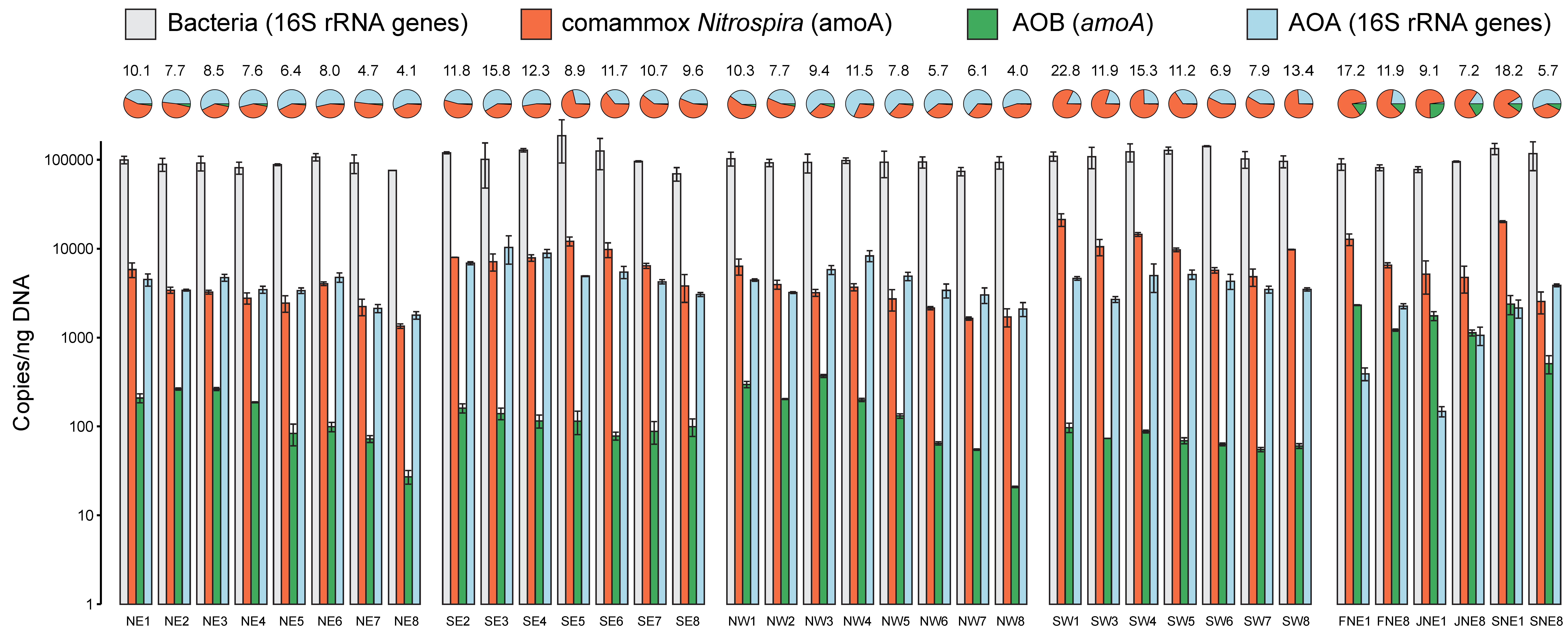

**Figure S1** Bacterial 16S rRNA, comammox *Nitrospira* (*amoA*)*,* AOB (*amoA*), and thaumarcheotal 16S rRNA gene abundances for all rotating biological contactor (RBC) samples. Error bars indicate standard deviation of qPCR technical duplicates. Pie charts show gene abundances as a proportion of all ammonia oxidizing prokaryotes. Numbers above the pie charts indicate the proportion of the total community that all ammonia oxidizing prokaryotes represent, as a percent. Gene copies were calculated based on the amount of DNA present in the original extractions but were not standardized to account for gene copy number. NE, northeast train; NW, northwest train; SE, southeast train; SW, southwest train.

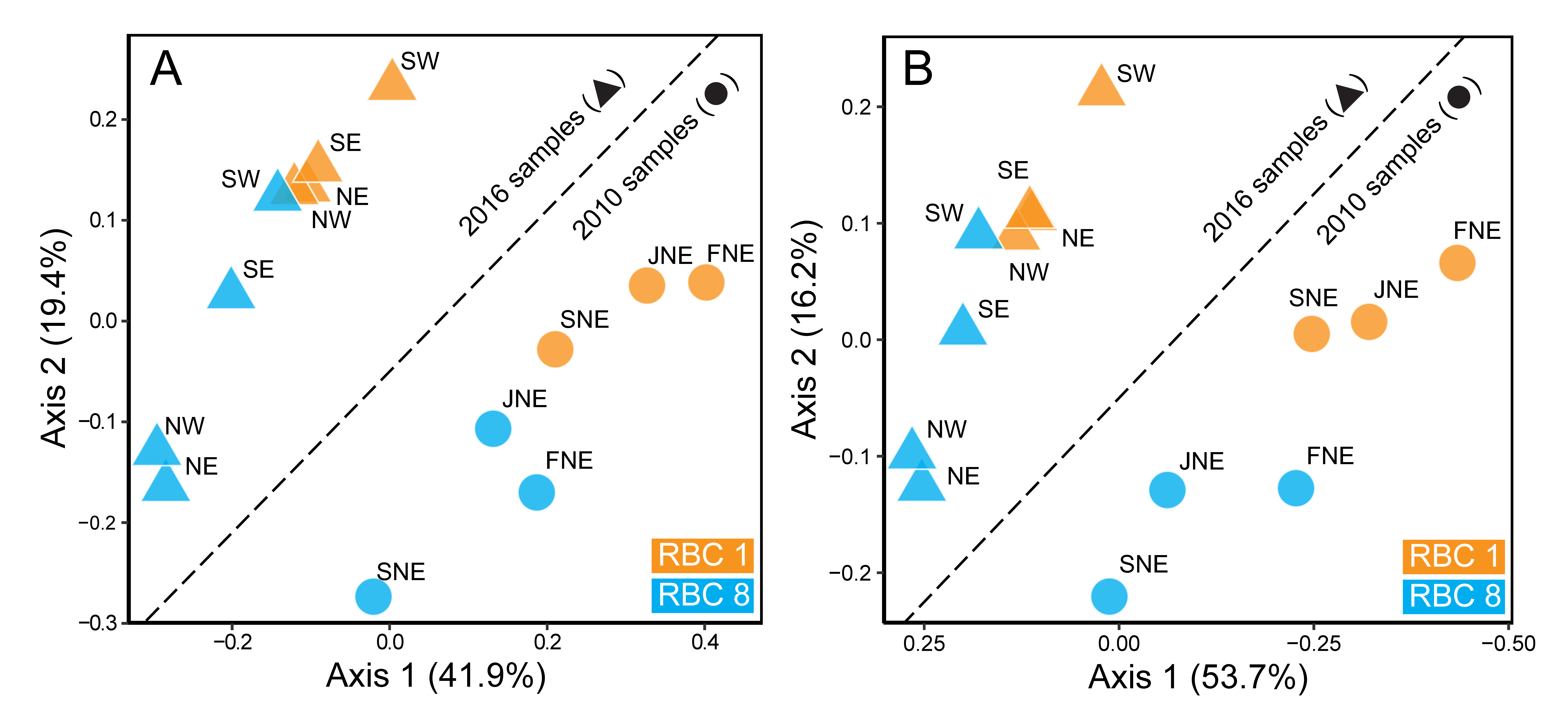

**Figure S2** Principal coordinate analysis (PCoA) using the Bray-Curtis dissimilarity metric for all 14 rotating biological contactor (RBC) samples analysed by 16S rRNA gene sequencing (A) and metagenome sequencing (B). The SE sample from 2016 was from RBC 2 instead of RBC 1, because SE1 was not operational at the time of sampling. NE, northeast train; NW, northwest train; SE, southeast train; SW, southwest train; F, February; J, June; S, September.

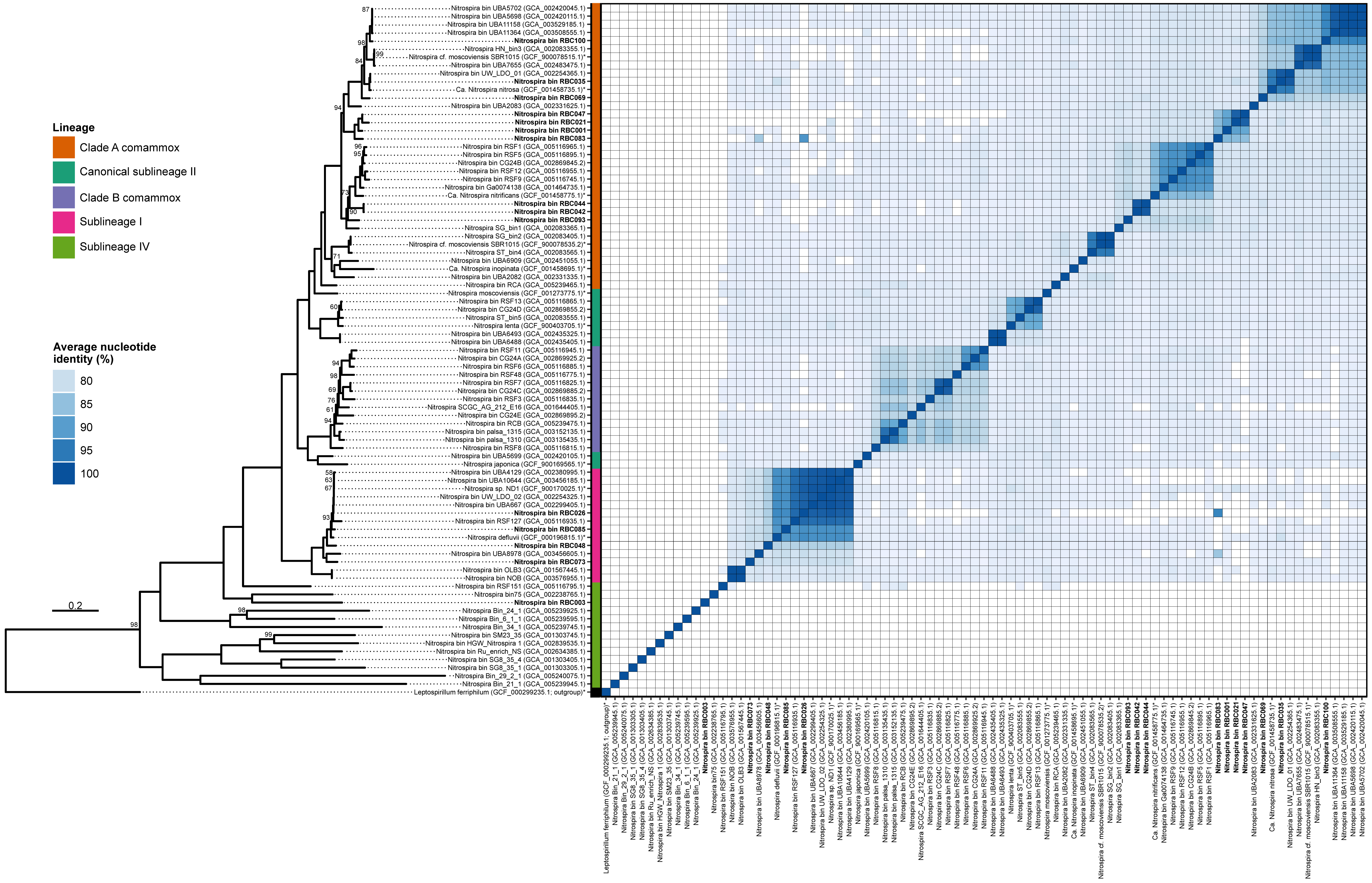

**Figure S3** Phylogeny and average nucleotide identity (ANI) of *Nitrospira* bins to reference genomes. The bins recovered from this study are indicated in bold, an asterisk indicates that the genome is from an enrichment or pure culture. The phylogeny was determined using a concatenated set of 74 core bacterial proteins, and ANI was determined using FastANI. All bootstrap values in the phylogeny are 100% except those shown, and the scale bar represents the proportion of amino acid change

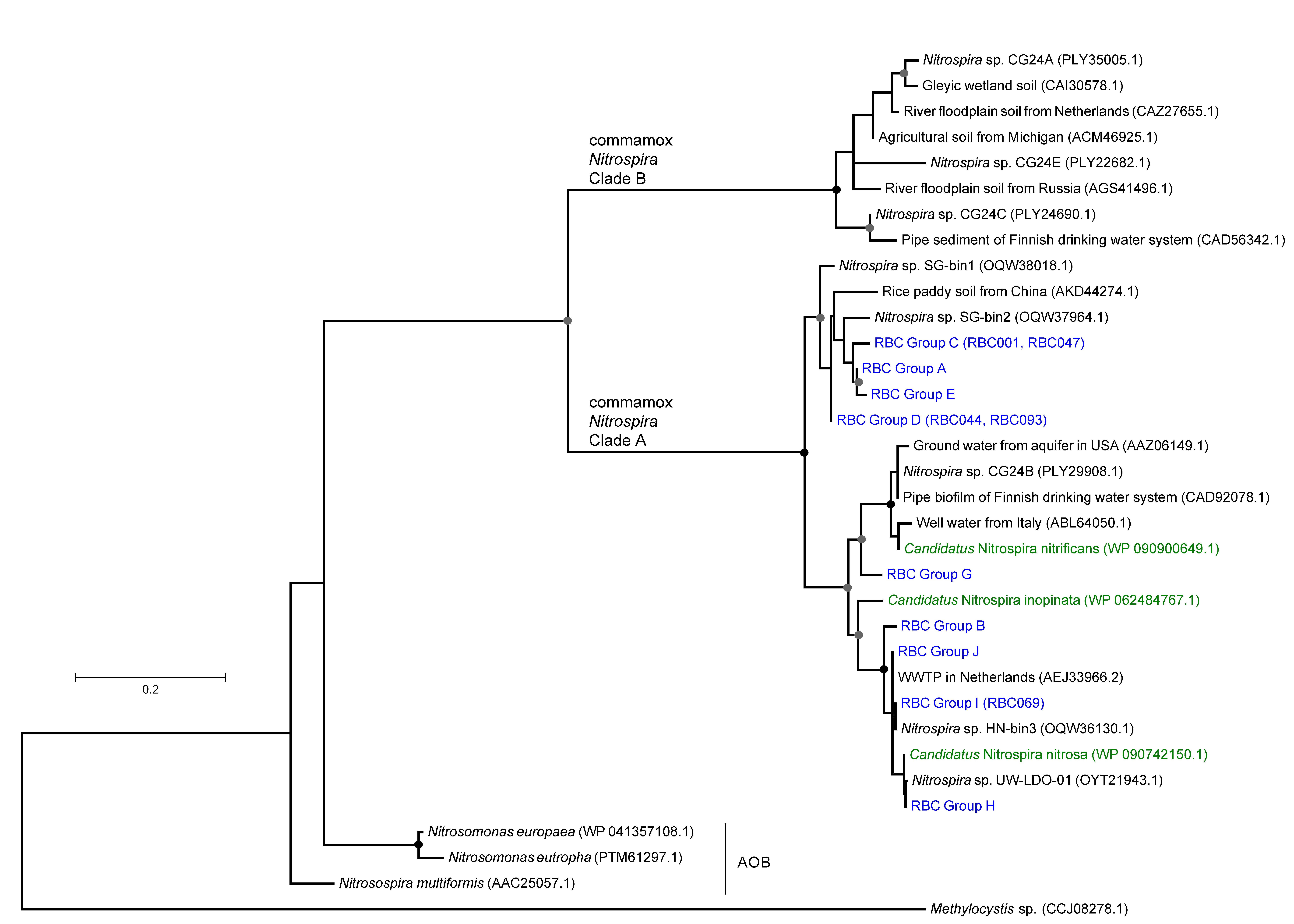

**Figure S4** Phylogeny of *Nitrospira* AmoA sequences.

Phylogenetic analysis of *Nitrospira* AmoA from assembled contigs was inferred using maximum likelihood analysis. Bootstrap values ≥ 60% are indicated with a grey dot, and those ≥ 90% are shown with a black dot. The tree is drawn to scale, with branch lengths measured in the number of amino acid substitutions per site. AmoA sequence groups from rotating biological contactors (RBCs) are shown in blue, with bins that encode the *amoA* sequences are indicated in brackets. Comammox *Nitrospira* species from enrichment cultures are shown in green. Comammox *Nitrospira* AmoA sequences from environmental surveys are shown in black. *Methylocystis* sp. PmoA was used as an outgroup.

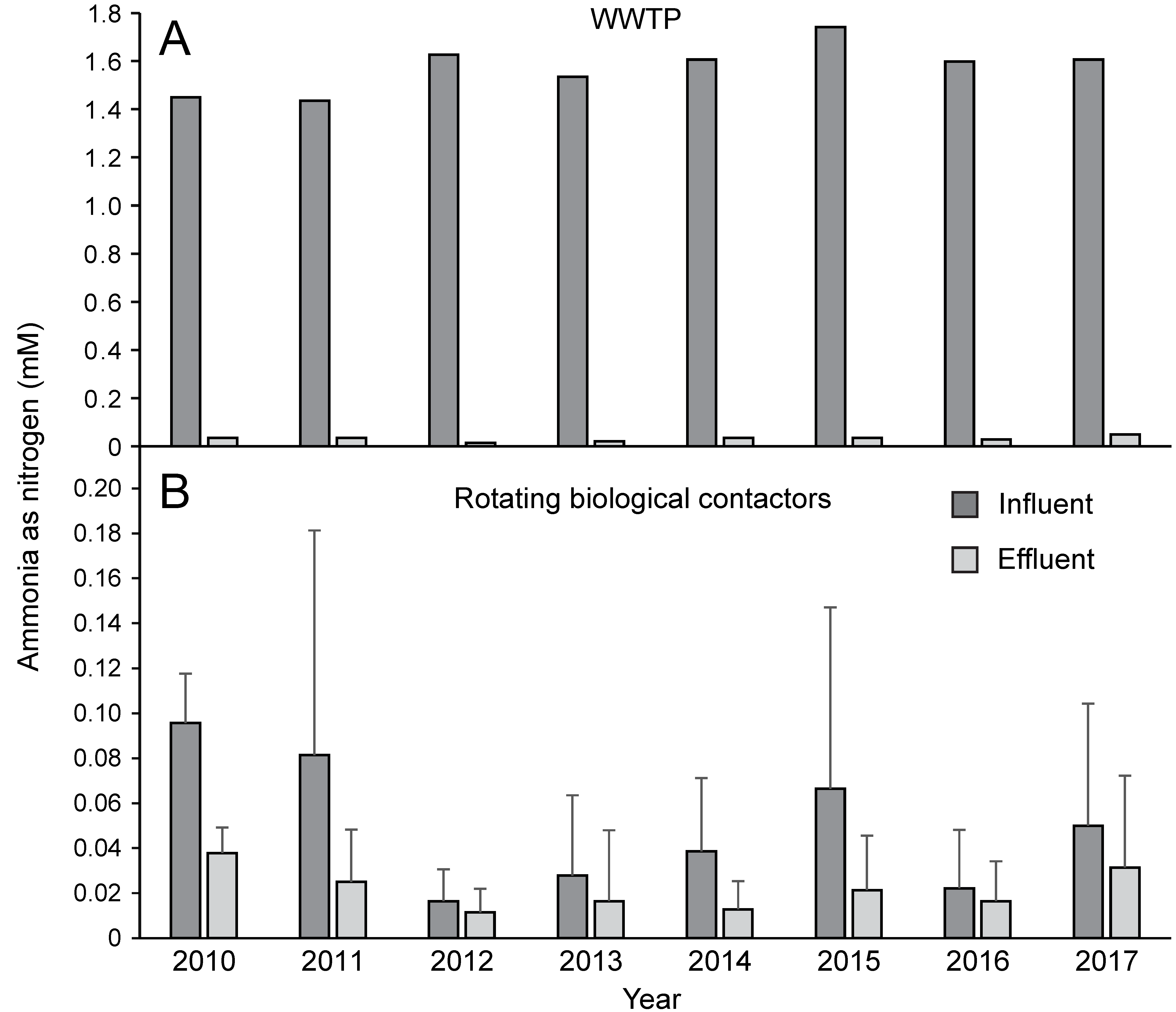

**Figure S5** Ammonia concentrations for the Guelph WWTP, obtained from WWTP operators. (A) Yearly average ammonia concentrations for Guelph WWTP influent (raw influent wastewater) and effluent (final effluent).

Values were obtained from the Guelph WWTP annual reports available online. (B) Average ammonia concentrations for rotating biological contactor (RBC) influent and effluent. The RBC influent values were measured as the secondary clarifier effluent combined, and the RBC effluent values were measured at the end of the southeast train. Values obtained from WWTP operators were for 24 hour composite samples, and were provided as mg/L values, and then converted to mM values. Averages for each month were calculated, and then averaged for the year. Error bars indicate standard deviation of yearly averages. Data for 2010 only included two months of data.
